# Cellular iron governs the host response to malaria

**DOI:** 10.1101/2023.02.05.527208

**Authors:** Sarah K. Wideman, Joe N. Frost, Felix C. Richter, Caitlin Naylor, José M. Lopes, Nicole Viveiros, Megan R. Teh, Alexandra E. Preston, Natasha White, Shamsideen Yusuf, Simon J. Draper, Andrew E. Armitage, Tiago L. Duarte, Hal Drakesmith

**Affiliations:** MRC Human Immunology Unit, MRC Weatherall Institute of Molecular Medicine, University of Oxford, John Radcliffe Hospital, Oxford, United Kingdom; Kennedy Institute of Rheumatology, Roosevelt Drive, OX3 7FY, Oxford, United Kingdom; Faculty of Medicine (FMUP) and Institute of Molecular Pathology, Immunology (IPATIMUP), University of Porto, Porto, Portugal; Instituto de Biologia Molecular e Celular & Instituto de Investigação e Inovação em Saúde (i3S), University of Porto, Porto, Portugal; Department of Biochemistry, University of Oxford, South Parks Road, Oxford, OX1 3QU, UK

**Author notes:** Authors contributed equally (Joe N. Frost, Felix C. Richter).

## Abstract

Malaria and iron deficiency are major global health problems with extensive epidemiological overlap. Iron deficiency-induced anaemia can protect the host from malaria by limiting parasite growth. On the other hand, iron deficiency can significantly disrupt immune cell function. However, the impact of host cell iron scarcity beyond anaemia remains elusive in malaria. To address this, we employed a transgenic mouse model carrying a mutation in the transferrin receptor (*Tfrc^Y20H/Y20H^*), which limits the ability of cells to internalise iron from plasma. At homeostasis *Tfrc^Y20H/Y20H^* mice appear healthy and are not anaemic. However, *Tfrc^Y20H/Y20H^* mice infected with *Plasmodium chabaudi chabaudi AS* showed significantly higher peak parasitaemia and body weight loss. We found that *Tfrc^Y20H/Y20H^* mice displayed a similar trajectory of malaria-induced anaemia as wild-type mice, and elevated circulating iron did not increase peak parasitaemia. Instead, *P. chabaudi* infected *Tfrc^Y20H/Y20H^* mice had an impaired innate and adaptive immune response, marked by decreased cell proliferation and cytokine production.

Moreover, we demonstrated that these immune cell impairments were cell-intrinsic, as *ex vivo* iron supplementation fully recovered CD4 T cell and B cell function. Despite the inhibited immune response and increased parasitaemia, *Tfrc^Y20H/Y20H^* mice displayed mitigated liver damage, characterised by decreased parasite sequestration in the liver and an attenuated hepatic immune response. Together, these results show that host cell iron scarcity inhibits the immune response but prevents excessive hepatic tissue damage during malaria infection. These divergent effects shed light on the role of iron in the complex balance between protection and pathology in malaria.

## INTRODUCTION

Malaria is a major global health problem that causes significant morbidity and mortality worldwide (1). It is caused by *Plasmodium* species parasites, which have a complex life cycle and are transmitted between humans by *Anopheles* mosquitos. In the human host, multiple cycles of asexual parasite replication inside red blood cells (RBC) result in extensive RBC destruction, immune activation, and microvascular obstruction (2). This blood stage of infection gives rise to symptoms such as fever, chills, headache, and malaise. In severe cases, it can also cause life-threatening complications such as acute anaemia, coma, respiratory distress, and organ failure (2).

There is a complex relationship between host iron status and malaria. Iron is an essential micronutrient that is required by most living organisms to maintain physiological and biochemical processes, such as oxygen transport and storage, cellular metabolism, and reduction-oxidation reactions (3,4). Despite the importance of iron, iron deficiency is exceedingly common in humans, and iron deficiency anaemia is estimated to affect a sixth of the world’s population (5,6). In the context of human malaria infection, iron deficiency can decrease the risk of disease, severe disease, and mortality (7–9). The protective effect of iron deficiency is at least partly mediated by anaemia, as RBCs isolated from anaemic individuals are less amenable to malaria parasite growth (10).

Meanwhile, oral iron supplementation is a risk factor for malaria in areas with limited access to preventative measures and treatment (11,12). This effect can to some extent be explained by iron supplementation stimulating erythropoiesis and increasing the proportion of reticulocytes and young erythrocytes, which are preferred targets for invasion by *P. falciparum* parasites (10). Malaria and iron deficiency also often disproportionally affect the same populations (e.g. young children in the WHO African Region) (1,6), in part, because malaria causes iron deficiency (13).

Anaemia is the primary and most profound consequence of iron deficiency. However, iron deficiency can also have other negative impacts on human health. Immune cells with high proliferative and anabolic capacities appear to be particularly sensitive to iron deficiency. As such, decreased iron availability can impair the proliferation and maturation of lymphocytes and neutrophils (14–16). Neutrophils and macrophages also require iron for enzymes involved in microbial killing (16–19). In animal models of iron deficiency, lymphocyte function is severely impaired, and the immune response to immunisation and viral infection is inhibited (20,21). Similarly, iron deficiency decreases inflammation and improves outcomes in mouse models of autoimmune disease (22–25). In humans, associations between iron deficiency and attenuated responses to some vaccines have been observed (20,21,26–28). Moreover, patients with a rare mutation in transferrin receptor-1 (TfR1), the primary receptor for iron uptake in cells, present with lymphocyte dysfunction and combined immunodeficiency (29,30).

Controlling a malaria infection requires two distinct but complementary immune responses. An early cell-mediated response, primarily driven by interferon-γ (IFN-γ) producing CD4^+^ T cells, prevents uncontrolled exponential parasite growth (31–35). Meanwhile, a humoral response is required to prevent recrudescence and to clear the infection (36,37). Excessive production of pro-inflammatory immune cells and cytokines can lead to sepsis-like complications and cause collateral damage to tissues and organs (38,39). Thus, the pro-inflammatory anti-parasite response must be balanced by immunoregulatory and tissue-protective responses to prevent immunopathology (40–43).

Although it is known that host iron deficiency influences malaria infection, the mechanisms that affect host health or *Plasmodium* virulence remain largely unknown. In particular, the effects of iron deficiency aside from anaemia, have scarcely been explored. Moreover, any effects on malaria immunity have not been investigated beyond a few observational studies that found associations between iron deficiency and attenuated antibody responses to malaria in children (7,44,45).

In this study, we aspired to deepen our understanding of how malaria infection is affected by host iron deficiency. To this end, we employed a genetic mouse model of cellular iron deficiency based on a rare mutation in TfR1 (*Tfrc^Y20H/Y20H^*), which causes combined immunodeficiency in humans (29,30). We found that decreasing host cellular iron levels increased peak malaria parasitaemia in mice infected with *P. chabaudi*. While *P. chabaudi*-induced anaemia and RBC invasion remained unaffected, the immune response to *P. chabaudi* was drastically inhibited. Interestingly, mice with cellular iron deficiency also had attenuated *P. chabaudi*-induced liver damage, suggesting reduced immunopathology. Hence, host cellular iron deficiency attenuated the immune response to malaria, leading to increased pathogen burden and mitigated liver pathology.

## RESULTS

### Decreased cellular iron uptake increases *P. chabaudi* pathogen burden

To investigate the effects of cellular iron availability on the host’s response to malaria, we utilised a transgenic mouse with a mutation in the cellular iron transporter TfR1. The *Tfrc^Y20H/Y20H^* mutation decreases receptor internalisation by approximately 50%, resulting in decreased cellular iron uptake (29). The effects of the *Tfrc^Y20H/Y20H^* mutation in erythroid cells are minimised due to a STEAP3-mediated compensatory mechanism (29). At homeostasis, adult *Tfrc^Y20H/Y20H^* mice are healthy, normal-sized, and not anaemic (Figure S1A-B). However, they have microcytic RBCs, compensated for by an increase in RBCs (Figure S1C-D), and mildly suppressed liver and serum iron levels (Figure S1E-F).

*Tfrc^Y20H/Y20H^* and wild-type mice were infected with a recently mosquito-transmitted rodent malaria strain, *P. chabaudi chabaudi* AS, which constitutively expresses GFP (hereafter referred to as *P. chabaudi*) (46,47) (Figure 1A). Recently mosquito-transmitted parasites were used to mimic a natural infection more closely, as vector transmission is known to regulate *Plasmodium* virulence and alter the host’s immune response (48–50). Consequently, parasitaemia is expected to be significantly lower upon infection with recently mosquito-transmitted parasites, compared to infection with serially blood-passaged parasites that are more virulent (47,48).

**Figure 1:**
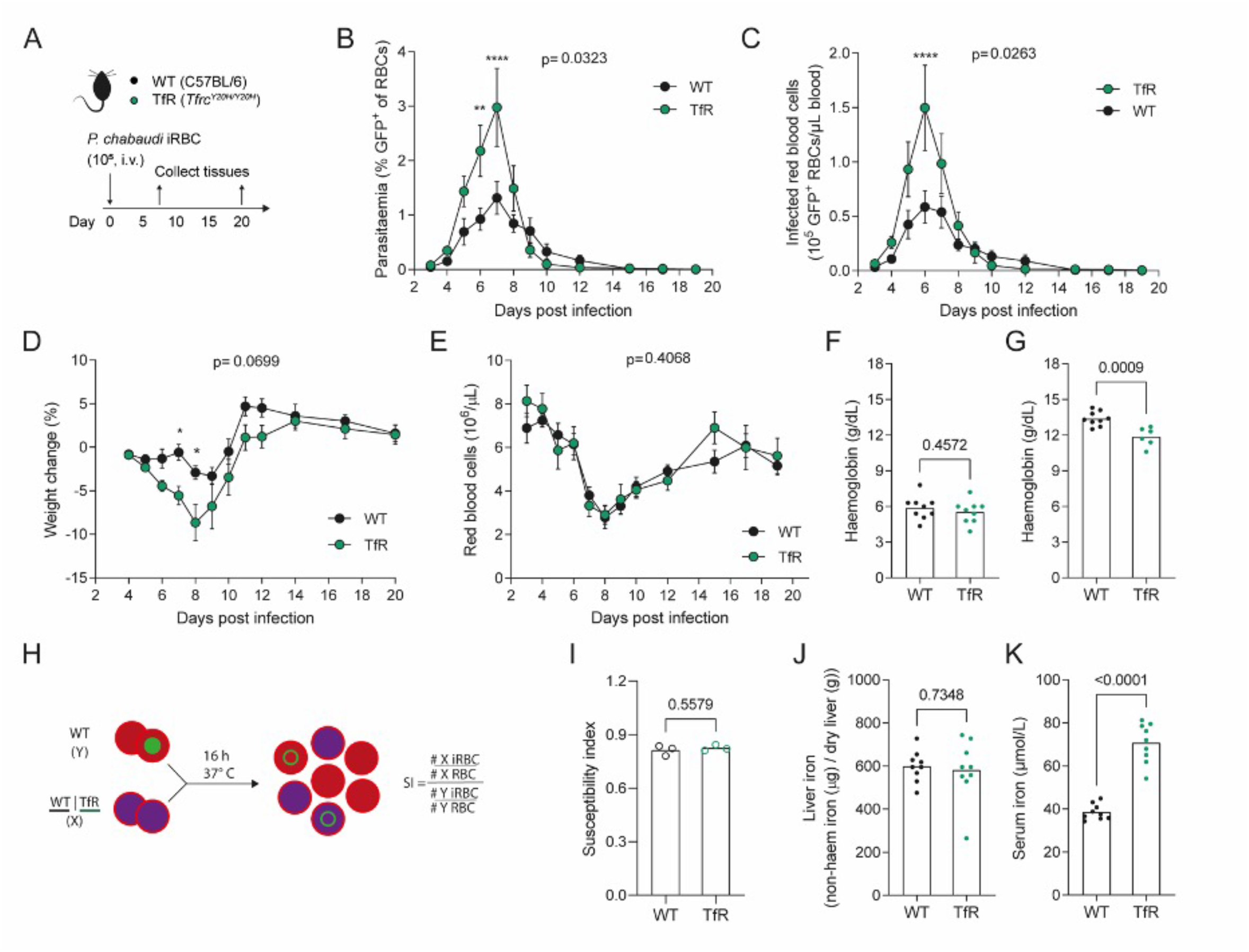
Decreased cellular iron uptake increases the *P. chabaudi* pathogen burden. **A)** C57BL/6 (WT) and *Tfrc^Y20H/Y20H^* (TfR) mice were infected by intravenous (i.v.) injection of 10^5^ recently mosquito-transmitted *P. chabaudi* infected red blood cells (iRBC). **B-E)** Parasitaemia (B), iRBC count (C), body weight change (D) and RBC count (E) measured throughout the course of infection. Mean ± SEM, mixed-effects analysis (B, C, E) or repeated measures two-way ANOVA (D), with Sidak’s multiple comparisons test, n=7-9. **F-G)** Haemoglobin measured 8 (F) and 20 (G) days after infection. Welch’s t-test, n=6-9. **H-I)** A mix of unlabelled WT RBC and iRBC were incubated with fluorescently labelled WT or TfR RBC and the invasion susceptibility index (SI) was determined after completion of a new invasion cycle. Mean, Welch’s t-test, n=3. **J-K)** Liver iron and serum iron levels measured 8 days after infection. Mean, Welch’s t-test, n=9.

Strikingly, mice with decreased cellular iron uptake had significantly higher peak parasitaemia and higher peak infected red blood cell (iRBC) counts (Figure 1B-C). The higher pathogen burden coincided with more severe weight loss than wild-type mice (Figure 1D). This phenotype contrasts previous studies, in which nutritional iron deficiency resulted in lower parasitaemia and increased survival of malaria infected mice (49,50). Hence, our findings highlight a distinct role for cellular iron in malaria pathology, which acts inversely to the protective effect of anaemia. This prompted us to investigate the cause of the higher parasite burden observed in our model.

### *Tfrc^Y20H/Y20H^* and wild-type mice have comparable malaria-induced RBC loss and anaemia

Anaemia-associated alterations of RBC physiology can affect malaria infection and have been put forward as the main cause of both the protective effect of iron deficiency and the increased risk associated with iron supplementation (10). We therefore monitored RBCs in wild-type and *Tfrc^Y20H/Y20H^*mice infected with *P. chabaudi.* Both genotypes displayed similar levels of malaria-induced RBC loss and RBC recovery (Figure 1E). Moreover, *Tfrc^Y20H/Y20H^* and wild-type mice were equally severely anaemic at the nadir of RBC loss, eight days post infection (dpi) (Figure 1F). At the chronic stage of infection (20 dpi), however, wild-type mice showed improved recovery from anaemia compared to *Tfrc^Y20H/Y20H^* mice (Figure 1G), consistent with a decreased ability of the *Tfrc^Y20H/Y20H^* cells to incorporate iron.

While anaemia and RBC counts were comparable between both genotypes during infection, it was nevertheless possible that differences in RBC physiology could alter the course of infection. Consequently, we performed an *in vitro* invasion assay to determine whether *Tfrc^Y20H/Y20H^* RBCs were more susceptible to *P. chabaudi* invasion. Fluorescently labelled wild-type or *Tfrc^Y20H/Y20H^* RBCs were incubated *in vitro* with RBCs from a *P. chabaudi* infected wild-type mouse. Upon completion of one asexual replication cycle, invasion was assessed, and the susceptibility index was calculated (Figure 1H). The RBC susceptibility indices of both genotypes were comparable (Figure 1I), thus indicating that the higher parasite burden in *Tfrc^Y20H/Y20H^* mice was not due to a higher susceptibility of their RBCs to *P. chabaudi* invasion.

### Hyperferremia does not substantially alter *P. chabaudi* infection

In addition to anaemia, it has been suggested that that variations in host iron levels could affect blood-stage *Plasmodium* parasite growth (51,52). Consequently, non-haem liver iron and serum iron was measured in wild-type and *Tfrc^Y20H/Y20H^* mice upon *P. chabaudi* infection. At the peak of infection, both genotypes had elevated liver and serum iron levels compared to homeostasis (Figure S1E-F & Figure 1J-K). Infected wild-type and *Tfrc^Y20H/Y20H^* mice had equivalent liver iron levels (Figure 1J), but serum iron levels were higher in *Tfrc^Y20H/Y20H^*mice (Figure 1K).

The elevated serum iron observed in infected *Tfrc^Y20H/Y20H^* mice was consistent with their restricted capacity to take up circulating transferrin-bound iron into tissues. However, we decided to investigate whether this supraphysiological serum iron (i.e., hyperferremia) could alter *P. chabaudi* parasite growth. To do this, we treated wild-type mice with a recombinant monoclonal anti-BMP6 IgG antibody (αBMP6) or an isotype control (Figure S2A). αBMP6 treatment suppresses hepcidin expression and elevates serum iron, as a consequence of unregulated release of iron from cellular stores (53) (Figure S2A). *P. chabaudi* infected mice treated with αBMP6 had higher serum iron than isotype control-treated mice on days 9 and 21 after infection (Figure S2B). Nevertheless, mice treated with αBMP6 and isotype had comparable peak parasitaemia and peak iRBC counts, although αBMP6 treated mice appeared to clear the parasites slightly more efficiently (Figure S2C-D). In addition, αBMP6 treatment did not significantly alter weight loss (Figure S2E). Taken together, this data indicates that hyperferremia, as observed in infected *Tfrc^Y20H/Y20H^* mice, does not increase peak parasitaemia. Accordingly, these findings further indicate that iron uptake by non-erythropoietic cells is decisive in the host response to malaria.

### Decreased cellular iron uptake attenuates the immune response to *P. chabaudi*

The immune response to malaria exerts control of parasitaemia, and the spleen is the main site of the immune response to blood-stage malaria (39,54). Therefore, we assessed the splenic immune response to *P. chabaudi* during the acute stage of infection (8 dpi). Interestingly, *Tfrc^Y20H/Y20H^* mice had attenuated splenomegaly during acute *P. chabaudi* infection (Figure 2A-B), suggesting a disrupted splenic response.

**Figure 2:**
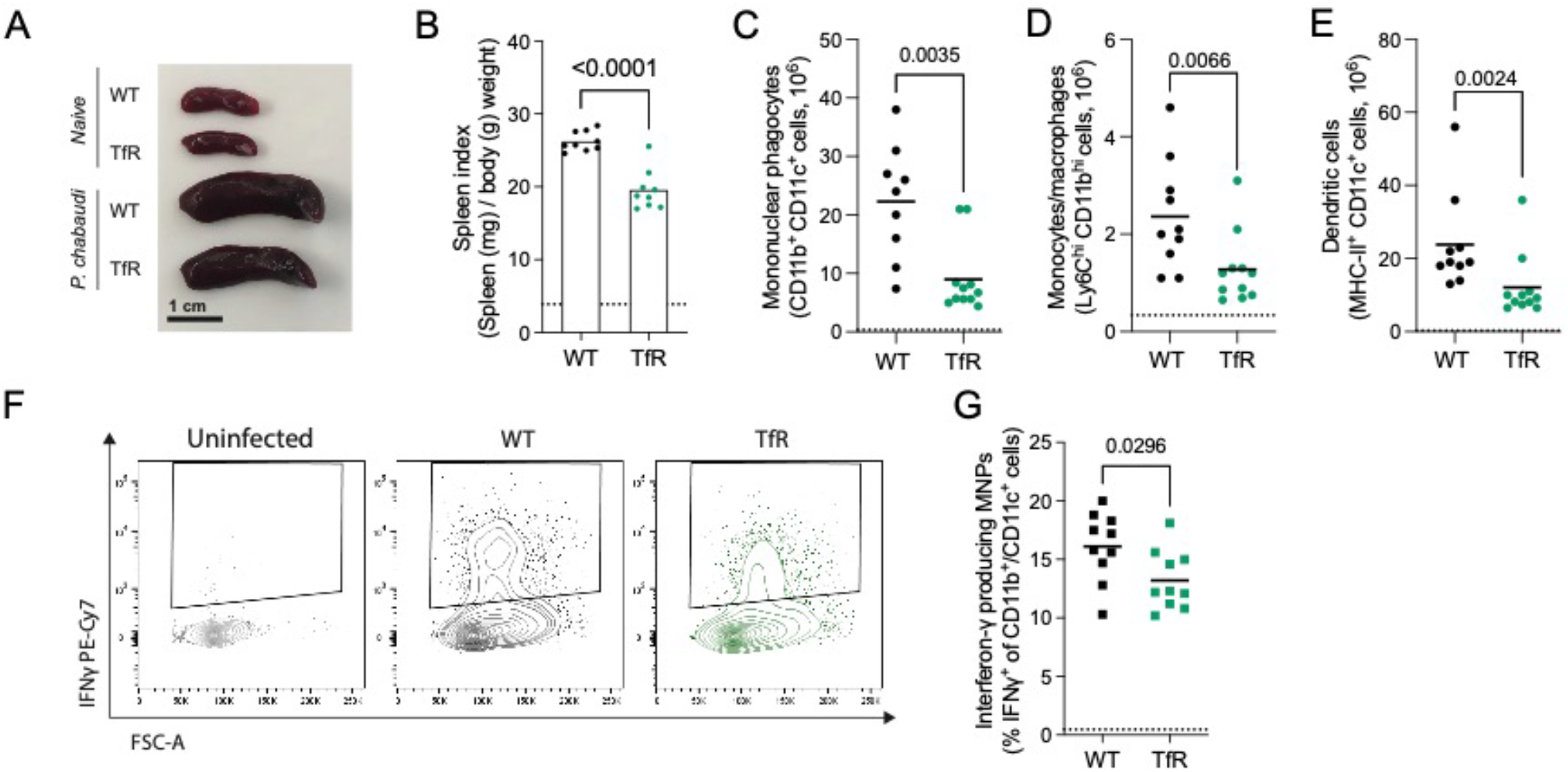
Decreased cellular iron uptake impairs the splenic MNP response to *P. chabaudi*. Splenic immune response to *P.chabaudi* in C57BL/6 (WT) and *Tfrc^Y20H/Y20H^* (TfR) mice at 8 dpi. **A)** Representative picture of spleens from naïve and *P. chabaudi* infected mice. **B)** Spleen index of spleens from naïve and *P. chabaudi* infected mice. Mean, Welch’s t-test n=9. **C-E)** Absolute numbers of CD11b^+^ CD11c^+^ mononuclear phagocytes (MNPs) (C), Ly6C^hi^ CD11b+ monocytes/macrophages (D) and MHCII^+^ CD11c^+^ dendritic cells (E) in spleens from naïve and *P. chabaudi* infected mice. Mean, Welch’s t-test on untransformed (C) or log transformed data (D, E) n=9-11. **F)** Representative flow cytometry plot of interferon-γ (IFNγ) production of CD11b^+^ CD11c^+^ MNPs. **G)** Proportion of IFNγ-producing MNPs, detected by intracellular cytokine staining. Mean, Welch’s t-test n=9-11. Dotted line represents uninfected mice.

Malaria infection leads to an influx of mononuclear phagocytes (MNP) into the spleen, where they are involved in cytokine production, antigen presentation, and phagocytosis of iRBCs (34,35,43). Upon *P. chabaudi* infection, fewer MNPs were detected in the spleen of *Tfrc^Y20H/Y20H^*mice (Figure 2C). This applied both to CD11b^+^ Ly6C^+^ MNPs (resembling inflammatory monocytes and/or monocyte-derived macrophages) and to CD11c^+^ MHCII^+^ MNPs (resembling dendritic cells) (Figure 2D-E & Figure S3A). In malaria infection, some MNPs can produce IFNγ that facilitates naïve CD4^+^ T cell activation and polarisation (34). Consequently, splenocytes from infected mice were cultured *ex vivo* with a protein transport inhibitor, and intracellular cytokine staining was performed. Interestingly, fewer MNPs from *Tfrc^Y20H/Y20H^* mice produced IFNγ compared to MNPs from wild-type mice (Figure 2F-G). Infected wild-type and *Tfrc^Y20H/Y20H^* mice had comparable splenic neutrophil, eosinophil and NK cell numbers during acute infection (8 dpi) (Figure S3B-D). Thus, mice with decreased cellular iron uptake had an attenuated MNP response to *P. chabaudi* infection.

### Cellular iron deficiency impairs the CD4^+^ T cell response to *P. chabaudi*

T cells, particularly CD4^+^ T cells, are a critical component of the immune response to blood-stage malaria (55). Therefore, we assessed the splenic T cell response to acute *P. chabaudi* infection. The total splenic CD4^+^ T cell count was comparable in both genotypes eight days after infection (Figure 3A). However, mice with decreased cellular iron uptake had a decreased proportion of effector CD4^+^ T cells (Figure 3B), and, consequently, fewer total splenic effector CD4^+^ T cells than wild-type mice (Figure 3C). In addition, the proportion of antigen-experienced CD44^+^ and PD1^+^ CD4^+^ T cells was also reduced in *Tfrc^Y20H/Y20H^* mice, re-enforcing their less activated state (Figure 3D-E). Moreover, fewer *Tfrc^Y20H/Y20H^*CD4^+^ T cells were actively dividing, based on the proliferation marker KI-67 (Figure 3F). This suggests a functional impairment of the CD4^+^ T cell response to *P. chabaudi* in mice with decreased cellular iron uptake.

**Figure 3:**
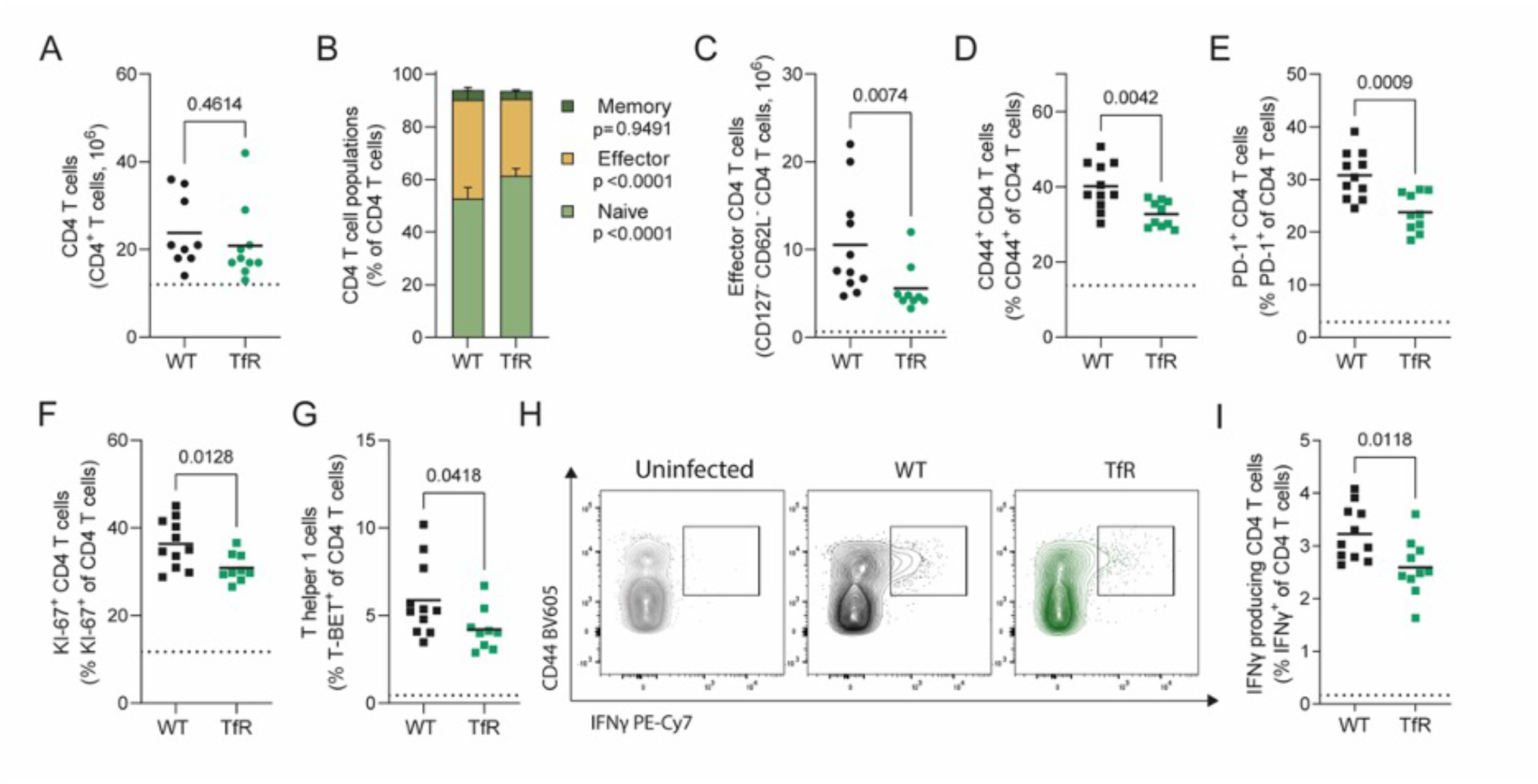
Decreased cellular iron uptake disrupts the effector CD4^+^ T cell response to *P. chabaudi*. Conventional CD4^+^ T cells (FOXP3^-^) in the spleen of *P. chabaudi* infected C57BL/6 (WT) and *Tfrc^Y20H/Y20H^* (TfR) mice, 8 dpi. **A)** Absolute number of CD4^+^ T cells in the spleen of *P. chabaudi* infected WT and TfR mice. Mean, Welch’s t-test, n=9-11. **B)** Proportions of naïve (CD44^-^ CD62L^+^), effector (CD62L^-^ CD127^-^) and memory (CD44^+^ CD127^+^) CD4^+^ T cells in the spleen of *P. chabaudi* infected WT and TfR mice. Mean, two-way ANOVA with Sidak’s multiple comparisons test, n=9-11. **C)** Absolute number of effector CD4^+^ T cells in the spleen of *P. chabaudi* infected WT and TfR mice. Mean, Mann-Whitney test, n=9-11. **D-E)** Proportions of CD4^+^ T cells expressing markers of antigen experience CD44^+^ (D) and PD-1^+^ (E) in the spleen of *P. chabaudi* infected WT and TfR mice. Mean, Welch’s t-test n=9-11 **F)** Proportion of proliferating (KI-67^+^) CD4^+^ T cells in the spleen of *P. chabaudi* infected WT and TfR mice. Mean, Welch’s t-test n=9-11 **G)** Proportion of T helper 1 (TBET^+^) CD4^+^ T cells in the spleen of *P. chabaudi* infected WT and TfR mice. Mean, Welch’s t-test n=9-11 **H)** Representative flow cytometry plot of IFNγ producing CD4^+^ T cells in the spleen of *P. chabaudi* infected WT and TfR mice. **I)** Proportion of IFNγ producing CD4^+^ T cells, detected by intracellular cytokine staining, in the spleen of *P. chabaudi* infected WT and TfR mice. Mean, Welch’s t-test n=10-11. Dotted line represents uninfected mice.

Similarly, the total CD8^+^ T cell count did not differ between genotypes (Figure S4A), but *P. chabaudi* infected *Tfrc^Y20H/Y20H^* mice had fewer effector CD8^+^ T cells eight days after infection (Figure S4B-C). However, there was no difference in the percentage of antigen-experienced (CD44^+^ or PD-1^+^) (Figure S4D-E), proliferating (KI-67^+^) (Figure S4F) or IFNγ producing (Figure S4G) CD8^+^ T cells. Hence the CD8^+^ T cell response to *P. chabaudi* infection was also attenuated, albeit to a lesser degree than CD4^+^ T cells.

T helper 1 (Th1) cells and other T helper subsets that express IFNγ are particularly important for malaria immunity (55). Interestingly, the proportion of CD4^+^ T cells that expressed the Th1 transcription factor T-BET was lower in mice with decreased cellular iron uptake (Figure 3G). Furthermore, fewer CD4 T cells from *Tfrc^Y20H/Y20H^* mice produced IFNγ upon *ex vivo* restimulation (Figure 3H-I). Thus, further strengthening the evidence of functional CD4^+^ T cell impairment in *Tfrc^Y20H/Y20H^* mice during *P. chabaudi* infection.

To determine whether these impairments were T cell intrinsic and iron-dependent, we utilized naïve CD4^+^ T cells isolated from uninfected wild-type and *Tfrc^Y20H/Y20H^* mice. The cells were cultured *in vitro* under Th1 polarising conditions for four days, in standard or iron-supplemented culture media (Figure 4A). *Tfrc^Y20H/Y20H^* lymphocytes can acquire iron under conditions where transferrin is hyper-saturated and sufficient quantities of free iron are likely to be generated (29,56). Proliferation was significantly impaired in *Tfrc^Y20H/Y20H^* CD4^+^ T cells but could be rescued in a dose-dependent manner by iron supplementation (Figure 4B-C). In addition, very few *Tfrc^Y20H/Y20H^* CD4^+^ T cells cultured in standard media produced IFNγ. However, iron supplementation completely rescued IFNγ production (Figure 4D-F). Hence, the CD4^+^ T cell deficiencies observed in *Tfrc^Y20H/Y20H^* mice during *P. chabaudi* infection were replicated *in vitro* and could be rescued by iron supplementation. These observations confirm that host cell iron scarcity disrupts CD4^+^ T cell function, leading to an inhibited CD4^+^ T cell response to *P. chabaudi* infection.

**Figure 4.**
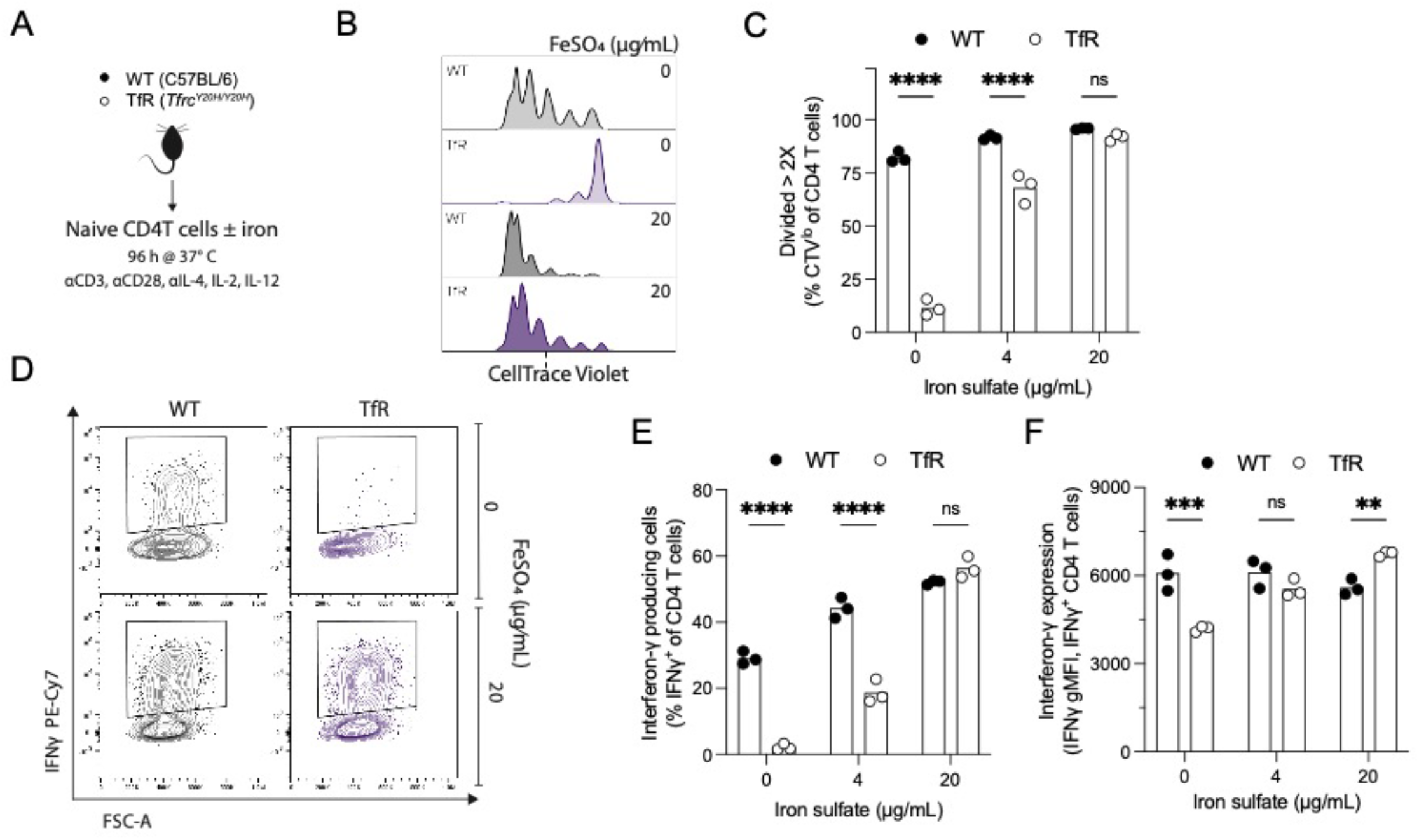
*In vitro* T helper 1 (Th1) polarised *Tfrc^Y20H/Y20H^* CD4^+^ T cells have impaired proliferation and effector function, which can be rescued by iron supplementation. **A)** Naïve CD4^+^ T cells were isolated from uninfected C57BL/6 (WT) and *Tfrc^Y20H/Y20H^* (TfR) mice, and cultured for 96 h in Th1 polarising media, with varying concentrations of iron sulfate (FeSO_4_). **B)**Representative flow cytometry plot of CD4^+^ T cell proliferation, quantified using CellTrace Violet. **C)** Proportion of CD4^+^ T cells that have divided more than two times (> 2X). Mean, two-way ANOVA, Sidak’s multiple comparisons test, n=3. **D)** Representative flow cytometry plot of IFNγ producing CD4^+^ T cell in the absence or presence of FeSO_4_. **E-F)** Proportion of IFNγ producing CD4 T cells (E) and IFNγ production per cell (F). Mean, two-way ANOVA, Sidak’s multiple comparisons test, n=3.

### Decreased cellular iron uptake disrupts the germinal centre response to *P. chabaudi*

An efficient germinal centre (GC) response is required to generate high-affinity antibodies that enable malaria clearance (36,37). In light of the impaired CD4^+^ T cell response to *P. chabaudi* in *Tfrc^Y20H/Y20H^* mice, we further examined the B cell supporting T follicular helper cell (Tfh) response. During the acute stage of infection, a smaller proportion of CD4^+^ T cells from *Tfrc^Y20H/Y20H^* mice expressed B cell co-stimulation receptor ICOS (Figure 5A). ICOS is essential in malaria infection, as it is required to maintain the Tfh cell response and sustain antibody production (57). In line with this, *Tfrc^Y20H/Y20H^* mice had fewer Tfh cells, both during the acute (8 dpi) and chronic (20 dpi) stages of infection (Figure 5B-C). Tfh cells support the activation, differentiation, and selection of high-affinity GC B cells, and are an essential component of the humoral immune response to malaria (37). Therefore, we next sought to assess the B cell response to *P. chabaudi* infection in *Tfrc^Y20H/Y20H^* and wild-type mice.

**Figure 5.**
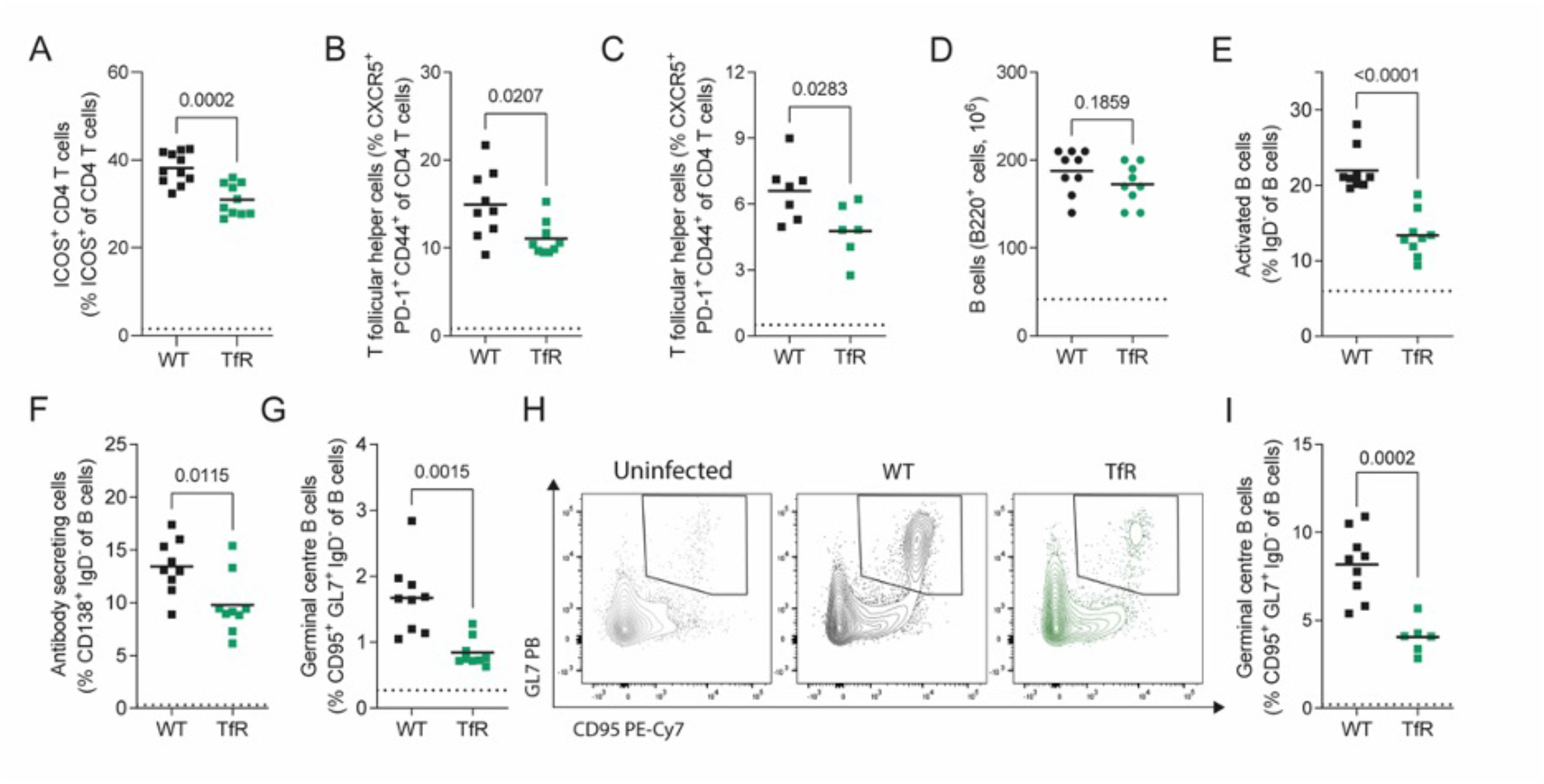
Decreased cellular iron uptake disrupts the germinal centre response to *P. chabaudi.* Splenic immune response of *P. chabaudi* infected C57BL/6 (WT) and *Tfrc^Y20H/Y20H^* (TfR) mice. **A)** Proportion of CD4^+^ T cells expressing B cell co-stimulatory receptor ICOS in the spleen of of *P. chabaudi* infected WT and TfR mice, 8 dpi. Mean, Welch’s t-test, n=10-11. **B)** Proportion of T follicular helper (Tfh) cells in the spleen of of *P. chabaudi* infected WT and TfR mice, 8 dpi. Mean, Welch’s t-test, n=9. **C)** Proportion of Tfh cells in the spleen of of *P. chabaudi* infected WT and TfR mice, 20 dpi. Mean, Welch’s t-test, n=6-7. **D-F)** Absolute total number of splenic B cells (D) and proportion of activated (E) and antibody secreting (F) splenic B cells in the spleen of of *P. chabaudi* infected WT and TfR mice, 8 dpi. Mean, Welch’s t-test, n=9. **G)** Proportion of germinal centre B cells in the spleen of of *P. chabaudi* infected WT and TfR mice, 8 dpi. Mean, Welch’s t-test, n=9. **H)** Representative flow cytometry plot of germinal centre B cells in the spleen of of *P. chabaudi* infected WT and TfR mice, 20 dpi. **I)** Proportion of germinal centre B cells in the spleen of of *P. chabaudi* infected WT and TfR mice, 20 dpi. Mean, Welch’s t-test on log transformed data, n=6-9. Dotted line represents uninfected mice.

We observed no difference between genotypes in the total number of splenic B cells at the acute stage of infection (8 dpi) (Figure 5D). However, mice with decreased cellular iron uptake had severely impaired B cell activation and fewer antibody-secreting effector B cells (Figure 5E-F). Additionally, *Tfrc^Y20H/Y20H^* mice had fewer GC B cells during acute infection (8 dpi) (Figure 5G). This effect remained in the chronic stage of infection (20 dpi) (Figure 5H-I), indicating a prolonged immune inhibition caused by restricted cellular iron availability.

### Cellular iron deficiency impairs B cell function

To determine if the *Tfrc^Y20H/Y20H^* mutation also had cell-intrinsic and iron-dependent effects on B cells, their functionality was further investigated *in vitro.* B cells were isolated from uninfected *Tfrc^Y20H/Y20H^*and wild-type mice, activated, and cultured in standard or iron-supplemented media for three days (Figure 6A). Expression of the B cell activation marker LAT-1 was lower on *Tfrc^Y20H/Y20H^* B cells than wild-type (Figure 6B). However, LAT-1 expression was rescued by iron supplementation, indicating improved B cell activation (Figure 6B). *Tfrc^Y20H/Y20H^* B cell proliferation was also severely impaired compared to wild-type cells, but was rescued by iron supplementation in a dose-dependent manner (Figure 6C-D). Iron scarcity also inhibited the potential of *Tfrc^Y20H/Y20H^* B cells to differentiate into antibody-secreting and class-switched cells (Figure 6E-G). This impairment was fully restored upon iron supplementation (Figure 6E-G). Overall, our data clearly show that the activation, proliferation, and differentiation of *Tfrc^Y20H/Y20H^* B cells were impaired, demonstrating that cellular iron deficiency causes cell-intrinsic B cell dysfunction.

**Figure 6.**
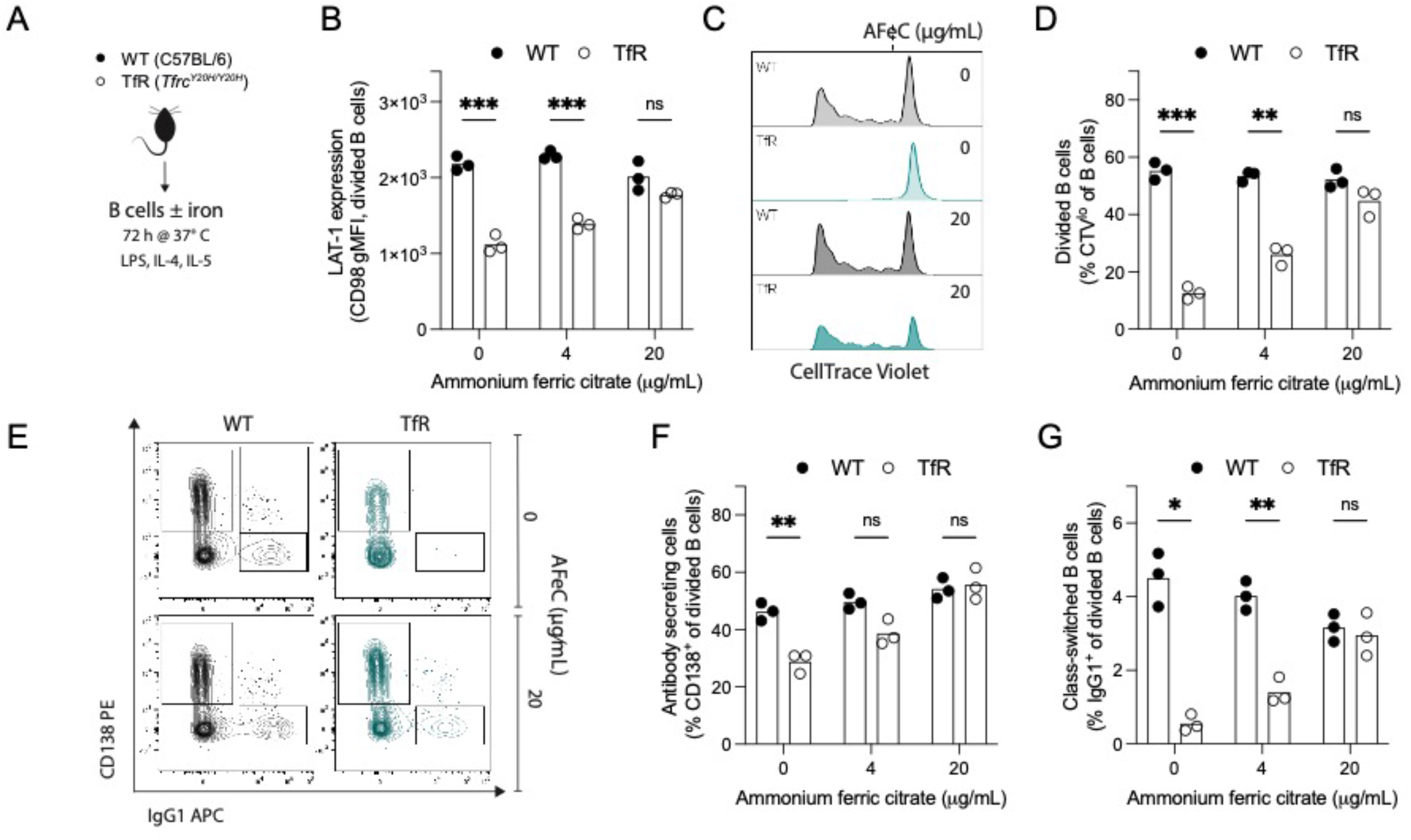
*In vitro* cultured *Tfrc^Y20H/Y20H^* B cells display impaired activation, proliferation and differentiation, which can be rescued by iron supplementation. **A)** B cells were isolated from uninfected C57BL/6 (WT) and *Tfrc^Y20H/Y20H^* (TfR) mice and cultured for 96 h in B cell activating media, with varying concentrations of ammonium ferric citrate (AFeC). **B)** Large neutral amino acid transporter-1 (LAT-1/CD98) expression on divided B cells. Mean, two-way ANOVA, Sidak’s multiple comparisons test, n=3. **C)** Representative flow cytometry plot of proliferating B cells, quantified using CellTrace Violet. **D)** Proportion of proliferating B cells (CTV^low^). Mean, two-way ANOVA, Sidak’s multiple comparisons test, n=3. **E)** Representative flow cytometry plots of antibody secreting (CD138^+^) and class-switched (IgG^+^) divided B cells. **F-G)** Proportion of antibody secreting (F) and class-switched (G) divided B cells. Mean, two-way ANOVA, Sidak’s multiple comparisons test, n=3.

### Decreased cellular iron uptake ameliorates *P. chabaudi*-induced liver pathology

*Tfrc^Y20H/Y20H^* mice experienced higher *P. chabaudi* parasitaemia and an inhibited immune response. However, the precise consequences of this disease phenotype remained unclear. Aspects of the immune response, such as the cytokine profile and the balance between pro-inflammatory and immunoregulatory responses, can tip the scales toward protection or pathology in malaria (39). Hence, an attenuated immune response could cause hyperparasitaemia, but it may also be crucial in limiting immunopathology. We therefore set out to characterise key indicators of malaria disease severity.

We first measured circulating levels of angiopoietin-2 (ANG-2) and alanine transferase (ALT). ANG-2 is a marker of endothelial activation that correlates with malaria disease severity and mortality in humans (58,59). Liver damage is also indicative of severe malaria (60), and ALT is a standard marker of liver damage. There was a trend towards lower ANG-2 and significantly decreased ALT in *Tfrc^Y20H/Y20H^* mice eight days after *P. chabaudi* infection, suggesting milder pathology (Figure 7A-B). Considering the substantial difference in serum ALT between genotypes, we further examined the malaria induced liver pathology. *Tfrc^Y20H/Y20H^* mice had lower expression of the tissue-damage and inflammation-induced acute phase protein genes *Saa1* and *Fga* (Figure S5A-B). Furthermore, while both genotypes developed malaria-induced hepatomegaly, there was a trend toward less severe hepatomegaly in *Tfrc^Y20H/Y20H^* mice (Figure S5C).

**Figure 7.**
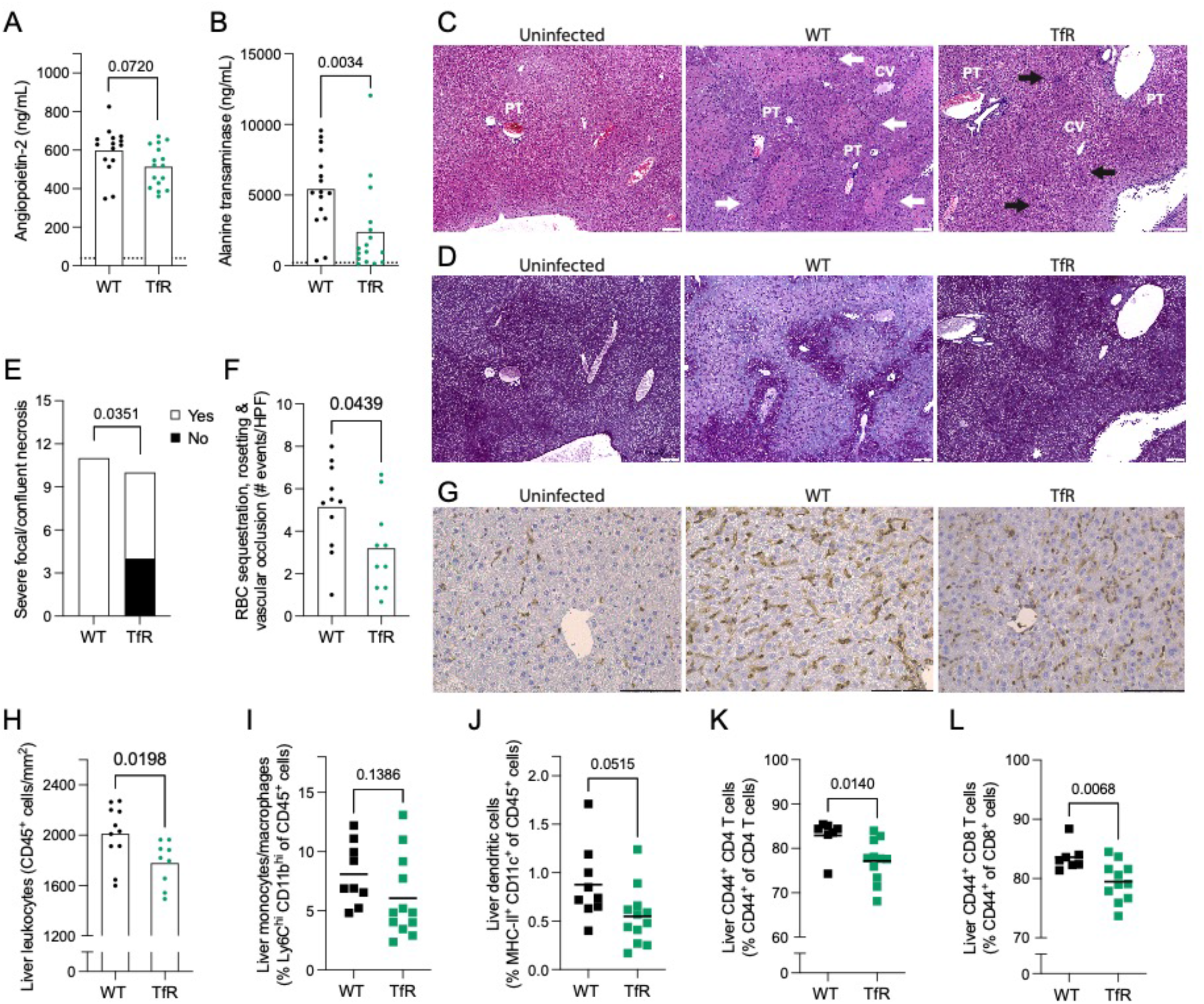
Decreased cellular iron uptake mitigates *P. chabaudi* liver pathology. Liver pathology of *P. chabaudi* infected C57BL/6 (WT) and *Tfrc^Y20H/Y20H^* (TfR) mice, 8 dpi. **A-B)** Serum levels of angiopoietin-2 (A) and alanine transaminase (B). Mean, Welch’s t-test, n=15-16. Dotted line represents uninfected mice **C-D)** Haematoxylin and eosin (C), and periodic acid–Schiff (D) staining of representative liver sections. Labels indicate central veins (CV), portal triads (PT), and areas of focal (black arrows) and bridging (white arrows) necrosis. Original magnification 40X, scale bar 100 µm. **E)** Quantification of severe hepatic necrosis (score ≧ 3) as measured by histological scoring. Count, Fisher’s exact test, n=10-11. **F)** Number of hepatic red blood cell sequestration, rosetting and vascular occlusion events per randomly imaged high-power field (HPF). Mean, Welch’s t-test, n=10-11. **G)** Immunohistochemistry staining of liver leukocytes (CD45+) in representative liver sections. Original magnification 20X, scale bar 100 µm. **H)** Quantification of CD45^+^ leukocytes in liver sections identified by immunohistochemistry staining. n= 9-11 **I-L)** Hepatic monocytes/macrophages (I), dendritic cells (J), CD44+ CD4+ T cells (K) and CD44+ CD8+ T cells (L). Mean, Welch’s t-test, n=7-12.

Histological analysis revealed hepatic pathology in all *P. chabaudi* infected mice, characterised by hepatocellular necrosis, sinusoidal dilatation, glycogen depletion, and infiltration by mononuclear immune cells (Figure 7C-D & Figure S5D-E). Interestingly, no polymorphonuclear immune cell infiltration was observed. All infected wild-type mice developed confluent necrosis (areas of lobular disarray, eosinophilia, and loss of glycogen deposits, score ≥3), and most individuals (8 out of 11) also displayed bridging necrosis (areas of confluent necrosis extending across multiple lobules, score=4) (Figure 7E & Figure S5F). In contrast, severe focal necrosis or confluent necrosis (score ≥3) was detected in just over half (6 out of 10) infected *Tfrc^Y20H/Y20H^* mice, and only four individuals developed bridging necrosis (Figure 7E & Figure S5F). Hence, the proportion of mice that developed severe hepatic necro-inflammation (score ≥3) upon *P. chabaudi* infection was significantly smaller in *Tfrc^Y20H/Y20H^* than in wild-type mice (Figure 7E).

Excess reactive liver iron and haem are known to cause liver damage in malaria (61,62). However, we observed no differences in total non-haem liver iron (Figure 1I) or liver lipid peroxidation, which correlates with ROS levels (Figure S5G). Hence, it is unlikely that tissue level variations in hepatic reactive iron or haem can explain the difference in liver damage. In addition, we measured the expression of two genes that are known to have a hepatoprotective effect in the context of iron loading in malaria: *Hmox1* (encodes haemoxygenase-1 (HO-1)) and *Fth1* (encodes ferritin heavy chain). Liver gene expression of *Hmox1* was higher in *Tfrc^Y20H/Y20H^* mice, while the expression of *Fth1* did not differ between genotypes, eight days after infection (Figure S5H-I). Thus, the higher expression of *Hmox1* may have contributed to a hepatoprotective effect in *Tfrc^Y20H/Y20H^* mice.

During malaria infection, endothelial activation leads to increased adhesion and sequestration of iRBCs, resulting in hepatic vascular occlusions and hypoxia that cause damage (2,63). Fewer sequestration, rosetting, and vascular occlusion events were detected in liver sections from *Tfrc^Y20H/Y20H^* mice eight days after *P. chabaudi* infection (Figure 7F). Together with the trend toward lower ANG-2 levels in *Tfrc^Y20H/Y20H^*mice (Figure 7A), this indicates that decreased endothelial activation and iRBC sequestration contributed to the attenuated liver pathology observed in *Tfrc^Y20H/Y20H^* mice.

Inflammation also causes severe disease and liver pathology in malaria (39,61,64). Hence, hepatic inflammation was approximated by measuring the expression of genes encoding pro-inflammatory cytokines IFNγ, TNFα, and IL-1β. We observed no difference in the expression of *Ifng* or *Tnf,* but *Il1b* expression was lower in *Tfrc^Y20H/Y20H^* mice eight days after *P. chabaudi* infection (Figure S5J-L). Moreover, immunohistochemistry staining showed reduced infiltration of leukocytes (CD45^+^ cells) in livers of *Tfrc^Y20H/Y20H^* mice (Figure 7G-H). Additionally, a smaller proportion of liver leukocytes (CD45^+^) were effector immune cells such as dendritic cells, CD44^+^ CD4^+^ T cells, and CD44^+^ CD8^+^ T cells (Figure 7I-L). Taken together, this data shows that host cell iron scarcity leads to an attenuated hepatic immune response during *P. chabaudi* infection.

## DISCUSSION

Iron deficiency impacts malaria infection in humans (7–9), but beyond the effects of anaemia (10), little is known about how host iron deficiency influences malaria infection. Here we investigated how restricted cellular iron acquisition influenced *P. chabaudi* infection in mice. *Tfrc^Y20H/Y20H^* mice developed comparable malaria-induced anaemia to wild-type mice, and RBC susceptibility to parasite invasion did not differ between genotypes. This therefore allowed us to largely decouple the effects of anaemia from other effects of iron on the host response to malaria. Strikingly, *Tfrc^Y20H/Y20H^* mice displayed an attenuated *P. chabaudi* induced splenic and hepatic immune response. This immune inhibition was associated with increased parasitaemia and mitigated liver pathology. Hence, for the first time, we show a role for host cellular iron acquisition via TfR1 in modulating the immune response to malaria, with downstream effects on both pathogen control and host fitness.

On first inspection, the higher parasite burden observed in *Tfrc^Y20H/Y20H^* mice may appear to be a severe consequence of cellular iron deficiency. In humans, however, high parasitaemia is not sufficient to cause severe disease (65). Moreover, the risk of severe malarial disease decreases significantly after only one or two exposures, whereas anti-parasite immunity is only acquired after numerous repeated exposures (2,66). It follows that mitigating immunopathology may be more important than restricting parasite growth for host survival. As previously noted, the *Tfrc^Y20H/Y20H^* mutation has relatively mild consequences for erythropoietic parameters compared to other haematopoietic lineages (29,30). However, in humans with normal TfR1-mediated iron uptake, iron deficiency sufficient to cause immune cell iron scarcity also normally causes anaemia (67). In such circumstances, parasite growth would likely be limited by anaemia, with the final result that iron deficiency may be protective overall, if it also minimises aspects of immunopathology.

Previous work has demonstrated the importance of regulating tissue haem and iron levels to prevent organ damage in malaria (61,62,68,69). For example, HO-1 plays an important role in detoxifying free haem that occurs as a result of haemolysis during malaria infection, thus preventing liver damage due to tissue iron overload, ROS and inflammation (61). Interestingly, infected *Tfrc^Y20H/Y20H^* mice had higher expression of *Hmox1*, but levels of liver iron and ROS comparable to that of wild-type mice. Consequently, this may be indicative of increased haem processing that could have a tissue protective effect. In humans, there is a correlation between transferrin saturation and ALT levels in patients with symptomatic malaria (62,70), suggesting that iron status may be linked to malaria-induced liver pathology in humans. However, it can be difficult to interpret measures of iron status in malaria infected individuals, since those parameters can be altered by inflammation and RBC destruction. Our findings reveal additional dimensions through which host iron status impacts malaria-induced tissue damage. The mitigated liver damage that we observed in *P. chabaudi* infected *Tfrc^Y20H/Y20H^* mice can likely be explained by a combination of factors; increased expression of hepatoprotective HO-1, decreased immune mediated endothelial activation, iRBC sequestration, and hepatic vascular occlusion, as well as, inhibited hepatic inflammation.

The pro-inflammatory immune response to malaria has downstream effects on cytoadherence, as pro-inflammatory cytokines activate endothelial cells, leading to higher expression of receptors for cytoadherence (2). As a consequence, *P. chabaudi* infected mice that lack adaptive immunity or IFNγ-receptor signalling, have substantially decreased sequestration of iRBCs in the liver, and no detectable liver damage (as measured by ALT) (63). Endothelial cells can also be activated by direct interactions with iRBCs (2), and in humans, ANG-2 correlates with estimated parasite biomass (59). However, although *P. chabaudi* infected *Tfrc^Y20H/Y20H^* mice had higher peak parasitaemia, they had fewer hepatic sequestration, rosetting, and vascular occlusion events and lower ANG-2 levels. The attenuated innate and adaptive immune response is the most probable cause of decreased endothelial activation and hepatic microvascular obstruction in *Tfrc^Y20H/Y20H^* mice. This, in turn, likely contributed to the clearly mitigated liver pathology, in spite of the higher parasitaemia. Upon *P. chabaudi* infection, we observed extensive infiltration of mononuclear leukocytes into the liver, but this response was repressed in *Tfrc^Y20H/Y20H^* mice. Specifically, infected *Tfrc^Y20H/Y20H^* mice had fewer effector-like immune cells in the liver. Hepatic immune cells can contribute to liver damage in malaria, for example, by producing pro-inflammatory cytokines or through bystander killing of hepatocytes (71). Consequently, a weaker hepatic pro-inflammatory immune response likely limited immunopathology and ameliorated malaria-induced liver damage in mice with cellular iron deficiency.

We have previously shown that hepcidin mediated hypoferremia inhibits the immune response to influenza infection in mice (21). In influenza, cellular iron scarcity exacerbated pulmonary tissue damage, because failed adaptive immunity led to an exacerbated inflammatory response and poor pathogen control (21). In contrast, we observed that decreased cellular iron acquisition inhibited both the innate and adaptive immune response to malaria, ultimately mitigating malaria-induced hepatic tissue damage and inflammation. This highlights the complex effects of iron deficiency on the immune system and underscores the need to consider its effect on different infectious diseases in a pathogen-specific manner. A better understanding of how host iron status affects immunity to infection could benefit the development of improved antimicrobial therapies and increase the safety of iron deficiency therapies.

The inhibited innate immune response to *P. chabaudi* in *Tfrc^Y20H/Y20H^* mice likely contributed to both the increased pathogen burden and the decreased liver pathology. Splenic MNPs are important for controlling parasitaemia (34,35,72), but MNPs are also vital for maintaining tissue homeostasis and preventing tissue damage in malaria (43,73). Although other innate cells, such as neutrophils, NK cells and γδT cells are an important part of the immune response to malaria, only the MNP response was distinctly impaired in *Tfrc^Y20H/Y20H^* mice. Notably, neutrophils are known to be sensitive to iron deficiency (16,74) and to affect both immunity and pathology in malaria (75,76). However, in the context of recently mosquito-transmitted *P. chabaudi* it appears that monocytes and macrophages, rather than granulocytes, may be particularly important for parasite control and tissue homeostasis (43,72).

CD4^+^ T cells and B cells become cell intrinsically dysfunctional during iron scarcity, as we have demonstrated *in vitro*. However, such cell-intrinsic effects are likely further aggravated by interactions with other iron-depleted cells *in vivo*. For example, CD4^+^ T cells support the B cell response to malaria (37,77), and the repressed CD4^+^ T cell response to *P. chabaudi* in *Tfrc^Y20H/Y20H^* mice presumably further constrained the B cell response. Proliferation is an aspect of immune cell function that appears to be particularly sensitive to iron deficiency (14,20,21). Unsurprisingly, we also see the most significant inhibitory effect on immune cell populations that expand greatly during *P. chabaudi* infection. In addition, proliferation is often required for lymphocyte differentiation and effector function (78), and the differentiation of Tfh and Th1 cells in malaria depends on a highly proliferative precursor CD4^+^ T cell subset (79). T cells from *Tfrc^Y20H/Y20H^* mice also had decreased KI-67 expression, further confirming impaired proliferation as a critical mechanism of immune inhibition under conditions of cellular iron scarcity. CD4^+^ T cells that produce pro-inflammatory cytokine are also sensitive to iron restriction, as we have shown for IFNγ, and as has been shown previously for IL-2 and IL-17 (80,81). Interestingly, iron overload can also alter CD4^+^ T cell cytokine production, and excess iron can have an inhibitory effect on IFNγ production (22,82). These observations underline that iron imbalance at either extreme can disturb immune cell function.

Despite the higher peak parasitaemia in *Tfr^cY20H/Y20H^* mice, both genotypes were able to clear *P. chabaudi* parasites at a comparable rate and prevent recrudescence. It follows that even a weakened humoral immune response appears to be sufficient to control *P. chabaudi* infection. However, our study did not investigate the effects of immune cell iron deficiency on the formation of long-term immunity, which may have been more severely affected. The impaired GC response, in particular, suggests that iron deficiency could counteract the formation of efficient immune memory to subsequent malaria infections. This is in line with human observational studies that have found a link between iron deficiency and weak antibody responses to *P. falciparum* (7,44,45). In humans, anti-parasite immunity forms very slowly and only after numerous repeated exposures to malaria infection (2). Some have suggested that this effect could be explained by impaired immune cell function in malaria (83,84), and future studies should consider whether inhibited immunity as a result of iron deficiency could contribute to this phenomenon. Moreover, the extensive geographical and epidemiological overlap of iron deficiency and malaria (1,6,13) makes this concept particularly relevant for further research.

It remains to be seen what the broader importance of cellular iron is in human malaria infection, in particular within the diverse genetic context of both humans and parasites, found in malaria endemic regions. Murine models of malaria are useful in providing hypothesis-generating results, but such findings ultimately ought to be confirmed and developed further through studies in human populations. This study revealed that decreased host cell iron acquisition inhibits the immune response to malaria and ameliorates hepatic damage, despite a higher parasite load and similar degree of anaemia, in mice. Altogether, our data highlight a previously underappreciated role for host cell iron in the trade-off between pathogen control and immunopathology, and add to our understanding of the complex interactions between iron deficiency and malaria. Hence, these findings have important implications for these two widespread and urgent global health problems.

## METHODS

### Mice

*Tfrc^Y20H/Y20H^* mice were initially provided by Professor Raif Geha, Boston Children’s Hospital/Harvard Medical School (29), and they were subsequently bred in-house at the University of Oxford. Control wild-type C57BL/6JOlaHsd mice were purchased from Envigo and co-housed with *Tfrc^Y20H/Y20H^* mice for 2-3 weeks prior to *P. chabaudi* infection. All mice were housed in individually ventilated specific-pathogen-free cages under normal light conditions (light 07.00-19.00, dark 19.00-07.00) and fed standard chow containing 188 ppm iron (SDS Dietex Services, diet 801161) ad-libitum. Age-matched, 8-13 week-old female mice were used for experiments. Females were exclusively utilised to prevent loss of animals due to fighting, and to minimise the risk of severe adverse events from *P. chabaudi* infection, which is higher in males (85). Euthanasia was performed through suffocation by rising CO_2_ concentrations, and death was confirmed by cervical dislocation.

### Ethics

All animal experiments were approved by the University of Oxford Animal Welfare and Ethical Review Board and performed following the U.K. Animals (Scientific Procedures) Act 1986, under project licence P5AC0E8C9.

### Parasites and infection

Transgenic recently mosquito-transmitted *P. chabaudi chabaudi* AS parasites expressing GFP (46,47) were obtained from the European Malaria Reagent Repository at the University of Edinburgh. To generate iRBCs for blood-stage *P. chabaudi* infections, frozen parasite stocks were rapidly thawed by hand and injected intraperitoneally (i.p.) into a single wild-type mouse. Once ascending parasitaemia reached 0.5-2%, the animal was euthanised and exsanguinated through cardiac puncture. Subsequent experimental infections were immediately initiated from the collected blood, by intravenously (i.v.) injecting 10^5^ iRBCs in 100 uL Alsever’s solution. Uninfected control mice received Alsever’s solution only.

To monitor *P. chabaudi* infection, blood was collected through micro-sampling from the tail vein of infected mice. Parasitaemia, iRBC count and RBC count was measured by flow cytometry, as previously described (46). Briefly, 2 μL of blood was diluted in 500 μL Alsever’s solution immediately after collection. The solution was further diluted 1:10 in PBS before acquisition on an Attune NxT Flow Cytometer (Thermo Fisher Scientific). A fixed volume of each sample was acquired, thus allowing for the enumeration of total RBCs and iRBCs per μL of blood.

### αBMP6 treatment

In order to experimentally raise serum iron levels, an αBMP6 human IgG monoclonal blocking antibody that cross-reacts with murine BMP6 (53) was administered. Control mice received a human IgG4 isotype control antibody. Both antibodies were diluted in 100 μL PBS and injected i.p at a dose of approximately 10 mg/kg body weight.

### Tissue processing

Organs and tissues were harvested shortly after euthanasia and kept cold until further analysis could be performed. Liver and spleen indices were calculated as the mass of the respective organs relative to mouse body weight. Blood was collected into appropriate blood collection tubes (BD Microtainer K2EDTA for whole blood or BD Microtainer SST/Sarstedt Microvette 100 Serum for serum), either by tail vein sampling or by cardiac puncture after euthanasia. Serum was prepared by centrifugation of the collection tubes at 10,000 x g for 5 min, and stored at −80° C.

### Blood analysis

RBC count, haemoglobin, and mean cell volume was measured from whole blood using an automatic KX-21N Haematology Analyser (Sysmex). Serum levels of ANG-2 and ALT were measured according to the producers’ instructions, using the Mouse ALT ELISA Kit (ab282882, Abcam) and the Mouse/Rat Angiopoietin-2 Quantikine ELISA Kit (MANG20, R&D Systems), respectively. Serum cytokines were measured using the LEGENDplex Mouse Inflammation Panel (740446, BioLegend) bead-based immunoassay. The assay was performed according to the manufacturer’s instructions, except that the protocol was adapted to use half-volumes.

### In vitro P. chabaudi invasion assay

To assess the susceptibility of wild-type and *Tfrc^Y20H/Y20H^* RBCs to *P. chabaudi* invasion, blood was collected from a *P. chabaudi* infected wild-type mouse during ascending parasitaemia (donor RBCs/Y), and from uninfected wild-type and *Tfrc^Y20H/Y20H^* mice (target RBCs/X). To remove leukocytes, the blood was passed through a cellulose (C6288, Merck) packed column, as previously described (86). The target RBCs were fluorescently labelled with 1 μM CellTrace Far Red (C34572, Thermo Fisher Scientific) in PBS, by diluting blood 1:10 with CellTrace solution and incubating in the dark for 15 min at 37° C, mixing the samples every 5 min. Afterward, the cells were washed twice in R10 media (RPMI-1640 with 10% FBS, 2 mM glutamine (G7513, Merck), 1% penicillin-streptomycin (P0781, Merck), 50 μM 2-Mercaptoethanol (31350, Thermo Fisher Scientific)) and resuspended in R10 media supplemented with 0.5 mM sodium pyruvate (1136007050, Thermo Fisher Scientific). 2 x 10^7^ donor RBCs and 2 x 10^7^ fluorescently labelled target RBCs were plated in the same well of a 96-well plate, and incubated overnight (∼16 h) in a candle jar at 37° C, to allow sufficient time for schizonts to develop and release merozoites. Invasion was measured as GFP^+^ RBCs and compared by calculating the susceptibility index, as previously described (87).

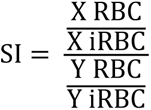

X = fluorescently labelled target wild-type or *Tfrc^Y20H/Y20H^* RBCs

Y = donor derived wild-type RBCs

### Iron measurements

Serum iron measurements were performed on an Abbott Architect c16000 automated analyser by Oxford University Hospitals Clinical Biochemistry staff using the MULTIGENT Iron Kit (Abbott), or using a Pentra C400 automated analyser with the Iron CP ABX Pentra Kit (HORIBA Medical).

Non-haem liver iron measurements were performed as previously described (88). In short, pieces of liver tissue were collected, snap-frozen, and stored at −80° C. The tissue was dried at 100° C for ∼6 h, weighed, and then digested in 10% trichloroacetic acid / 30% hydrochloric acid in water for ∼20 hours at 65°C. Subsequently, a chromogen reagent containing 0.1% bathophenanthrolinedisulphonic acid (Sigma, 146617) / 0.8% thioglycolic acid (Sigma, 88652) / 11% sodium acetate in water was added, and the absorbance at 535 nm measured. The iron content was determined by comparing the samples against a standard curve of serially diluted ammonium ferric citrate (F5879, Merck).

### Flow cytometry

Single cell suspensions for flow cytometry were prepared through mechanical and enzymatic dissociation. Spleens were passed through 70 μM cell strainers, incubated with 120 Kunitz U/mL deoxyribonuclease I (DN25, Merck) in R10 for 15 min with agitation, and passed through 40 μM cell strainers. Livers were perfused with PBS with 10% FBS prior to harvest. To prepare single cell suspensions, the livers were disrupted with scissors, incubated with 0.5 mg/mL collagenase IV (C5138, Merck) and 120 Kunitz U/mL DNAse I in R10 for 45 min with agitation, and passed through 70 μM cell strainers. RBC lysis was subsequently performed by resuspending pelleted cells in tris-buffered ammonium chloride buffer (0.017 M Tris / 0.14 M NH_4_Cl, adjusted to pH 7.2 with HCl) and incubating for ∼5 min on ice before washing with R10.

Immune cells were isolated from livers by Percoll (17-08-91, GE Healthcare) separation. Single-cell suspensions were gently overlayed onto 33% Percoll and centrifuged for 25 min at 800 x g. After centrifugation, the supernatant was discarded and the remaining leukocytes were washed twice with R10.

For intracellular cytokine staining, splenocytes were cultured *ex vivo* in R10 at 5-2 x 10^5^ cells/mL, in round-bottom tissue culture treated 96-well plates, with protein transport inhibitor Brefeldin A for 4-6 h at 37° C, 5% CO^2^. To activate T cells, 0.5 μg/mL anti-mouse CD3 (100201, BioLegend) was added to splenocytes from *P. chabaudi* infected mice.

Cells were counted using a CASY Cell Counter and Analyser (BOKE), and 1-5 x 10^6^ cells were stained for flow cytometry. The cells were washed in PBS, blocked with TruStain FcX (101319, BioLegend), and stained with a viability dye (NIR Fixable Viability Kit (42301/5, BioLegend) or LIVE/DEAD Fixable Near-IR Dead Cell Stain Kit (L34975, Thermo Fisher Scientific)) for ∼10 min at 4° C in the dark. Next, fluorophore-conjugated antibodies were added to the cells and incubated for ∼20 min. The cells were washed twice in PBS and fixed by incubating with Fixation Buffer (420801, BioLegend) for ∼10 min at 4° C in the dark. Alternatively, the cells were fixed and permeabilised using eBioscience FOXP3/Transcription Factor Staining Buffer Set (00-5523-00, Thermo Fisher Scientific), and transcription factor staining was performed, according to the manufacturer’s instructions. Intracellular cytokine staining was performed after permeabilization with Intracellular Staining Permeabilization Wash Buffer (421002, BioLegend) for ∼30 min, according to the manufacturer’s protocol. The samples were acquired on an Attune NxT or BD LSR Fortessa X-20 (BD) flow cytometer.

### *In vitro* culture of primary immune cells

Naïve CD4^+^ T cells and B cells were isolated according to the manufacturer’s instructions from mixed splenocyte and lymph node single-cell suspensions using the EasySep Mouse Naïve CD4^+^ T Cell Isolation Kit (19765, STEMCELL), or from splenocyte single-cell suspensions using the EasySep Mouse B Cell Isolation Kit (19854, STEMCELL). The isolated cells were fluorescently labelled with 5 μM CellTrace Violet (C34571, Thermo Fisher Scientific) in PBS for 8 min at 37° C and washed twice in R10 media. Cell counting was performed with a CASY Cell Counter and Analyser.

For CD4^+^ T cells, flat-bottom tissue culture treated 96-well plates were pre-coated with 5 μg/mL anti-mouse CD3 and the cells were seeded at 5 x 10^5^ cells/mL. They were cultured in Th1-polarising media consisting of R10 with 1 μg/mL anti-mouse CD28 (102101, BioLegend), 5 μg/mL anti-mouse IL-4 (504102, BioLegend), 10 ng/mL IL-12 (505201, BioLegend), 25 U/mL IL-2 (575404, BioLegend) and 50 μM 2-Mercaptoethanol. The media was replaced after 48 h of culture. To iron supplement the culture medium, iron sulphate heptahydrate (F8633, Merck) was added at the previously specified concentrations.

B cells were cultured at 7.5 x 10^5^ cells/mL in flat-bottom tissue culture treated 96-well plates, in R10 media with 1% MEM amino acids (11130, Thermo Fisher Scientific), 2 μg/mL LPS (tlrl-peklps, InvivoGen), 10 ng/mL IL-4 (574302, BioLegend), 10 ng/mL IL-5 (581502, BioLegend) and 50 μM 2-Mercaptoethanol. Ammonium ferric citrate was added at the specified concentrations to iron supplement the media.

CD4^+^ T cells were cultured for 96 h and B cells for 72 h at 37° C, 5% CO2, before flow cytometry staining. The type of iron used to supplement the culture media was chosen to optimise cell viability.

### Gene expression analysis

Gene expression analysis by quantitative real-time PCR, was performed on liver samples preserved in RNAlater Stabilization Solution and stored at −80° C (AM7020, Thermo Fisher Scientific). The tissue was homogenised with a TissueRuptor II (9002725, QIAGEN) before total RNA was extracted using the RNeasy Plus Mini Kit (74136, QIAGEN), according to the manufacturer’s protocols. cDNA was synthesised using the High-Capacity RNA-to-cDNA Kit (4387406, Thermo Fisher Scientific) and subsequent gene expression analysis was performed on 1-5 ng/mL cDNA, using TaqMan Gene Expression Master Mix (4369016, Thermo Fisher Scientific) and the TaqMan Gene Expression Assays (Thermo Fisher Scientific) listed in Table 1, all according to the manufacturers’ instructions. An Applied Biosystems 6500 Fast Real-Time PCR System (Thermo Fisher Scientific) instrument was used to run the samples, and the relative gene expression was calculated through the 2^-ΔCT^ method (89).

**Table 1.** List of TaqMan Gene Expression Assays.

| Protein | Gene | Assay code |
| --- | --- | --- |
| Fibrinogen alpha chain | *Fga* | Mm00802584_m1 |
| Hypoxanthine-guanine phosphoribosyltransferase | *Hprt* | Mm01545399_m1 |
| Interferon $\gamma$ | *Ifng* | Mm01168134_m1 |
| Interleukin 1β | *Il1b* | Mm00434228_m1 |
| Serum amyloid A1 | *Saa1* | Mm00656927_g1 |
| Tumour necrosis factor $\alpha$ | *Tnf* | Mm00443258_m1 |

### Liver histology

Liver samples were fixed with 4% paraformaldehyde in PBS and embedded in paraffin. Following deparaffinization with xylene and hydration by a passage through a grade of alcohols, 3 µm-thick sections were stained with haematoxylin-eosin, and Periodic Acid-Schiff, before and after diastase digestion, at IPATIMUP Diagnostics, Portugal, using standard procedures.

Histopathology scores for lobular necro-inflammatory activity were assigned using the criteria of Scheuer (90) for the grading of chronic hepatitis. In short, the scores were assigned as follows, 0 = inflammation absent, 1 = inflammation but no hepatocellular death, 2 = focal necrosis (one or a few necrotic hepatocytes/acidophil bodies), 3 = severe focal death, confluent necrosis without bridging, and 4 = damage includes bridging necrosis. Sections were scored independently by two investigators with experience in liver histopathology who were blinded to the experimental groups. The total numbers of RBC endothelial cytoadherence (sequestration), rosetting and vascular occlusion events were counted blindly in random high-power (×400 magnification) fields of liver sections. Images were captured using an Olympus BX50 photomicroscope.

For the immunohistochemical detection of CD45^+^ cells, liver sections were subjected to antigen retrieval with citrate buffer, endogenous peroxidases were blocked with 0.6% H_2_O_2_ and non-specific antigens were blocked with 5 % bovine serum albumin. Samples were incubated with goat anti-mouse CD45 antibody (1:50, AF114, R&D Systems, MN, USA) followed by horseradish peroxidase-conjugated rabbit anti-goat IgG (1:250, R-21459, ThermoFisher Scientific). Immunoreactivity was visualized using 3,3’-diaminobenzidine. Quantification was performed by counting positive cells in 5 random fields per liver at 200× magnification using QuPath Open Software for Bioimage Analysis (version 0.4.0).

### Thiobarbaturic acid reactive substances assay

Liver ROS/lipid peroxidation was appreciated by quantifying malondialdehyde, using the TBARS Assay Kit (700870, Cayman Chemical) as described by the manufacturer. Briefly, tissue homogenates were prepared from snap-frozen liver tissue by adding 1 mL RIPA buffer per 100 mg of tissue, and lysing using Precellys soft tissue homogenising tubes (KT03961-1-003.2, Bertin Instruments) according to manufacturers instruction. The lysates were allowed to react with thiobarbaturic acid at 95° C for 1 h, cooled on ice, and centrifuged for 10 min at 1,600 x g at 4° C. Subsequently, the absorbance of the lysates at 530 nm was measured.

### Software and statistical analysis

All flow cytometry data analysis was performed using FlowJo analysis software (BD). Graphs were generated using GraphPad Prism (GraphPad Software).

Statistical analysis was also performed in GraphPad Prism and differences were considered statistically different when p<0.05 (* p<0.05, ** p<0.01, *** p<0.001, **** p<0.0001). The D’Agostino-Pearson omnibus normality test was used to determine normality/lognormality. Parametric statistical tests (e.g. Welch’s t-test) were used for normally distributed data. For lognormal distributions, the data was log-transformed prior to statistical analysis. Where data did not have a normal or lognormal distribution, or too few data points were available for normality testing, a nonparametric test (e.g. Mann-Whitney test) was applied. A t-test (or a comparable nonparametric test) was used to compare the means of two groups. As a rule, t-tests were performed with Welch’s correction, as it corrects for unequal standard deviations but does not introduce error when standard deviations are equal. Two-way ANOVA was used for analysis with two categorical variables and one continuous variable. The applied statistical test and sample size (n) is indicated in each figure legend.

## ACKNOWLEDGEMENTS

The authors thank the staff of the University of Oxford Department of Biomedical Services for assistance with animal husbandry and procedures, and the Weatherall Institute of Molecular Medicine Flow Cytometry Facility for technical assistance. We are also grateful to the Clinical Biochemistry Unit (Oxford University Hospitals NHS Foundation Trust), and Samira Lakhal-Littleton and Goran Mohammad at the Oxford University Department of Physiology, Anatomy & Genetics for assistance with biochemical measurements. Additionally, we would like to sincerely thank Wiebke Nahrendorf, Philip Spence and Joanne Thompson at the University of Edinburgh for helpful scientific discussions and for providing us with *P. chabaudi* parasites.

## FUNDING

This work was supported by:

Wellcome Trust Infection, Immunology & Translational Medicine doctoral programme grant (awarded to SKW, grant no. 108869/Z/15/Z)

UK Medical Research Council (MRC Human Immunology Unit core funding awarded to HD, grant no. MC_UU_12010/10)

Portuguese National Funds through FCT—Fundação para a Ciência e a Tecnologia, I.P., under the project UIDB/04293/2020

Wellcome Trust Infection, Immunology & Translational Medicine doctoral programme grant (awarded to FCR, grant no. 203803/Z16/Z)

## CONFLICT OF INTERESTS

The authors declare that they have no conflict of interest.

---

**Figure S1:**
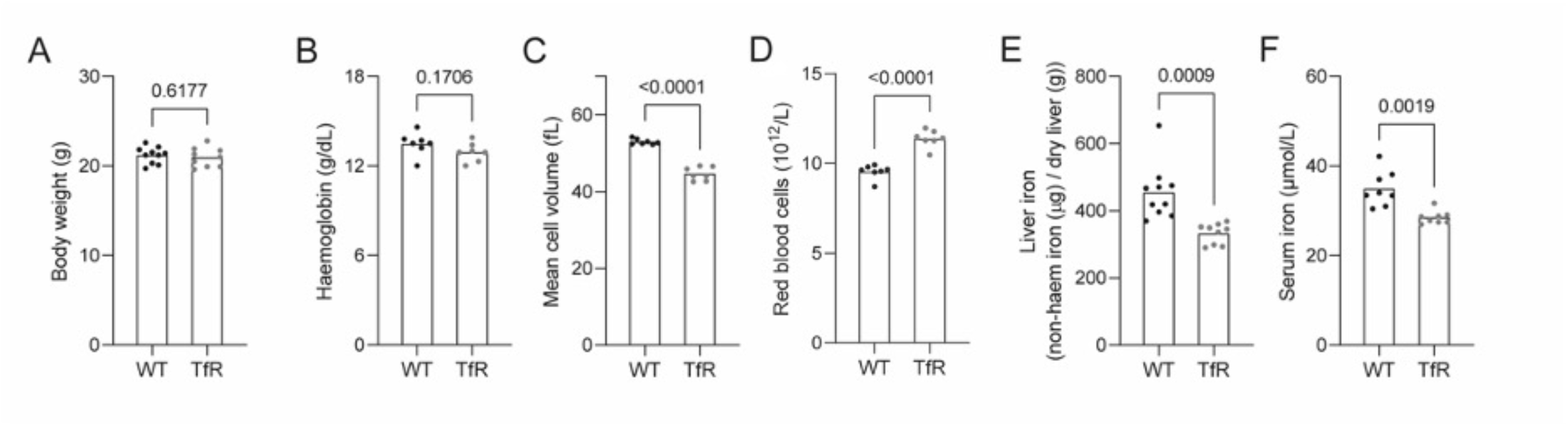
*Tfrc^Y20H/Y20H^* mice have mild microcytosis and decreased iron levels at homeostasis. Uninfected 8–12-week-old C57BL/6 (WT) and *Tfrc^Y20H/Y20H^* (TfR) mice were used for characterization. **A)** Body weight at homeostasis. Mean, Welch’s t-test, n=9-10. **B-D)** Haemoglobin (B), mean RBC volume (C) and RBC count (D) at homeostasis. Mean, Welch’s t-test, n=7. **E-F)** Liver iron (E) and serum iron (F) at homeostasis. Mean, Welch’s t-test, n=8-10.

**Figure S2.**
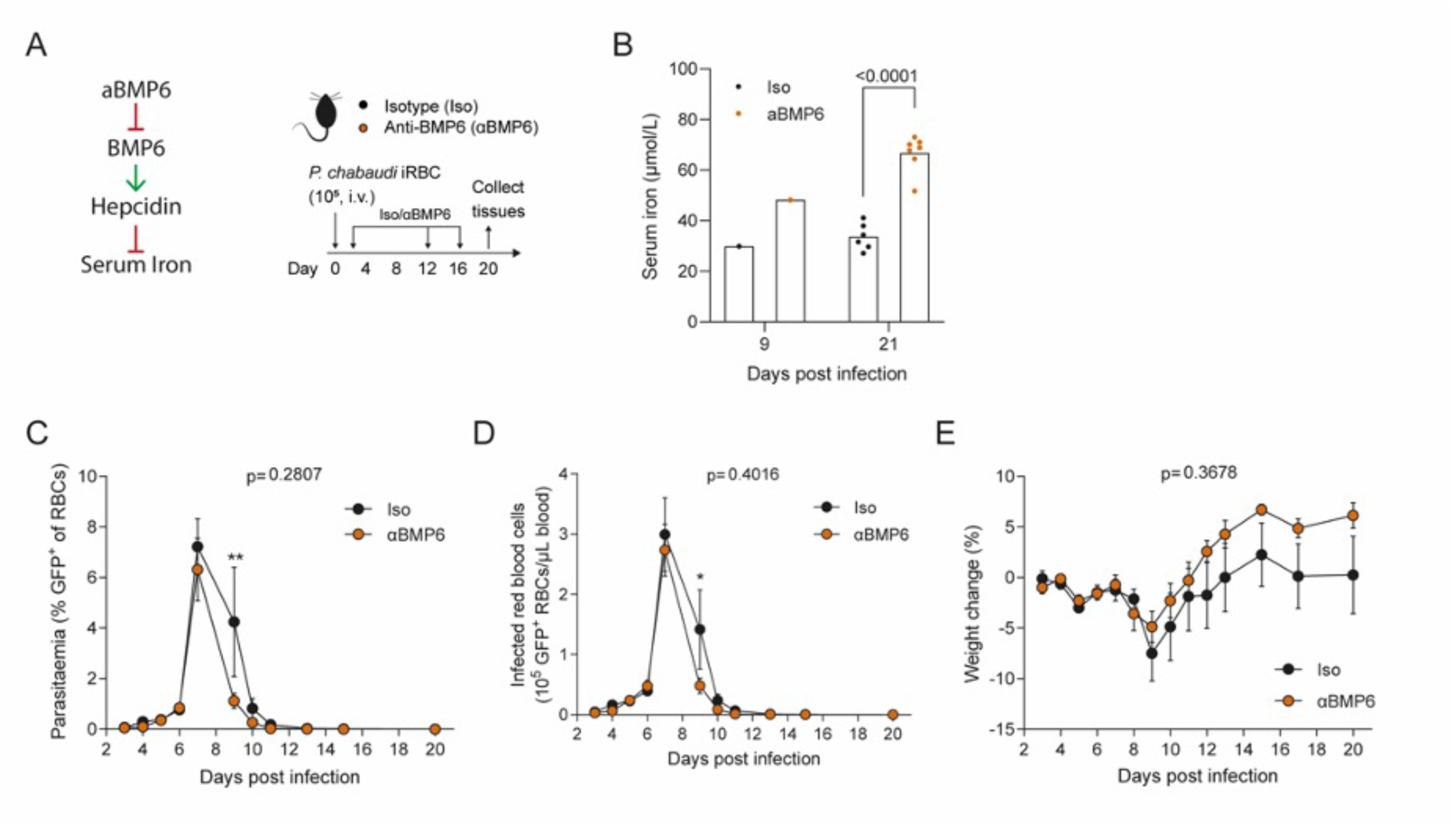
Hyperferremia does not increase *P. chabaudi* parasitaemia. **A)** C57BL/6 mice were infected by intravenous (i.v.) injection of 10^5^ *P. chabaudi* infected red blood cells (iRBC). A monoclonal anti-BMP-6 antibody (αBMP6) or an isotype control antibody (Iso) was administered 2, 12 and 16 days after infection. **B)** Serum iron measured 9 and 21 dpi in mice treated with anti-BMP6 or Iso. At day 9 post-infection, serum samples, collected through tail bleeding, were pooled for each experimental group to obtain sufficient sample for the quantification. At day 21 post-infection, mice were sacrificed, and serum samples collected through cardiac puncture. Mean, Welch’s t-test, n=6-8. **C-E)** Parasitaemia (C), iRBC count (D) and relative change in body weight (E) were measured throughout the course of infection. Mean ± SEM, two-way ANOVA with Sidak’s multiple comparisons test, n=6-8.

**Figure S3:**
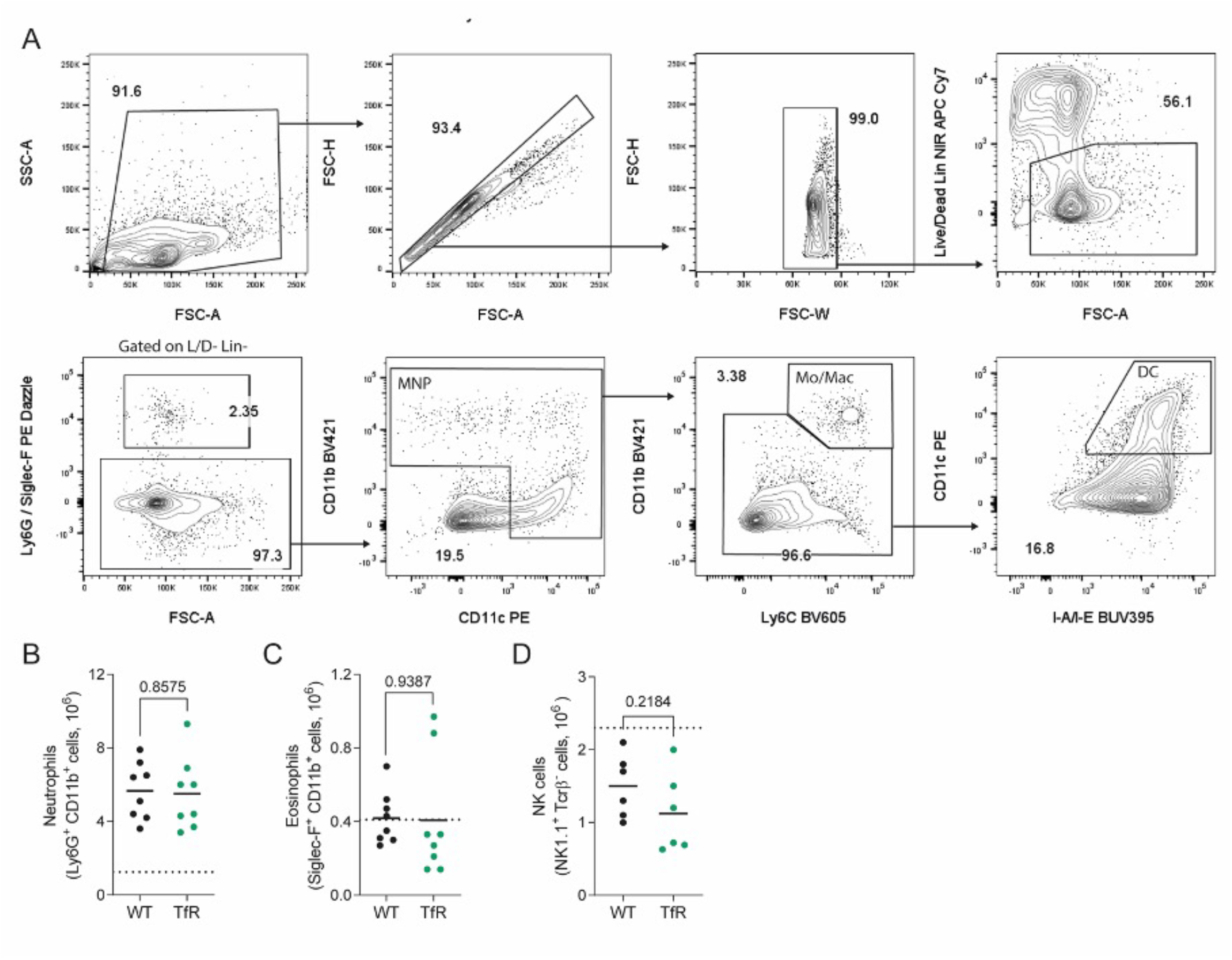
Mononuclear phagocyte gating scheme and innate immune response to *P. chabaudi* infection. Splenic immune response of *P. chabaudi* infected C57BL/6 (WT) and *Tfrc^Y20H/Y20H^* (TfR) mice, 8 dpi. **A)** Gating strategy for mononuclear phagocytes (MNP), monocytes/macrophages (Mo/Mac) and dendritic cells (DC). **B-D)** Absolute number of splenic neutrophils (B), eosinophils (C) and NK cells (D) of WT and TfR mice at 8dpi. Mean, Welch’s t-test, n = 6-8.

**Figure S4:**
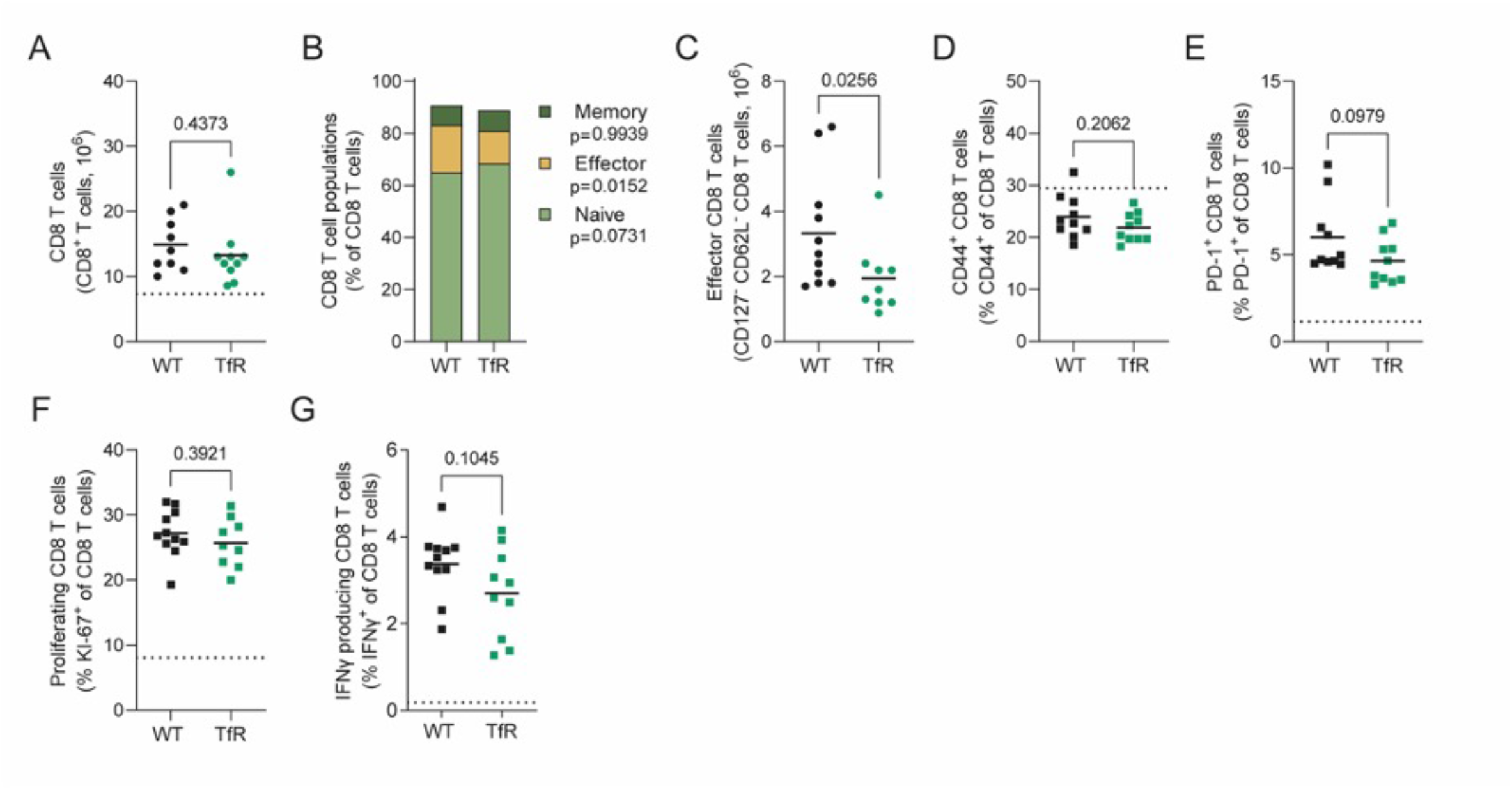
Decreased cellular iron uptake attenuates the effector CD8^+^ T cell response to *P. chabaudi*. CD8^+^ T cells in the spleen of *P. chabaudi* infected C57BL/6 (WT) and *Tfrc^Y20H/Y20H^* (TfR) mice, 8 dpi. **A)** Absolute numbers of splenic CD8^+^ T cells of *P. chabaudi* infected WT and TfR mice. Mean, Welch’s t-test, n=9-10. **B)** Proportion of naïve (CD44^-^ CD62L^+^), effector (CD62L^-^ CD127^-^) and memory (CD44^+^ CD127^+^) splenic CD8^+^ T cells of *P. chabaudi* infected WT and TfR mice. Mean, two-way ANOVA with Sidak’s multiple comparisons test, n=9-11. **C)** Absolute number of effector CD8^+^ T cells of spleens from *P. chabaudi* infected WT and TfR mice. Mean, Mann-Whitney test, n=9-11. **D-E)** Proportion of splenic CD8^+^ T cells expressing markers of antigen experience CD44^+^ (D) and PD-1^+^ (E) of *P. chabaudi* infected WT and TfR mice. Mean, Welch’s t-test n=10 **F)** Proportion of proliferating (KI-67^+^) splenic CD8^+^ T cells of *P. chabaudi* infected WT and TfR mice. Mean, Welch’s t-test n=9-11 **G)** Proportion of IFNγ producing splenic CD8^+^ T cells, detected by intracellular cytokine staining of *P. chabaudi* infected WT and TfR mice. Mean, Welch’s t-test n=10-11. Dotted line represents uninfected mice.

**Figure S5.**
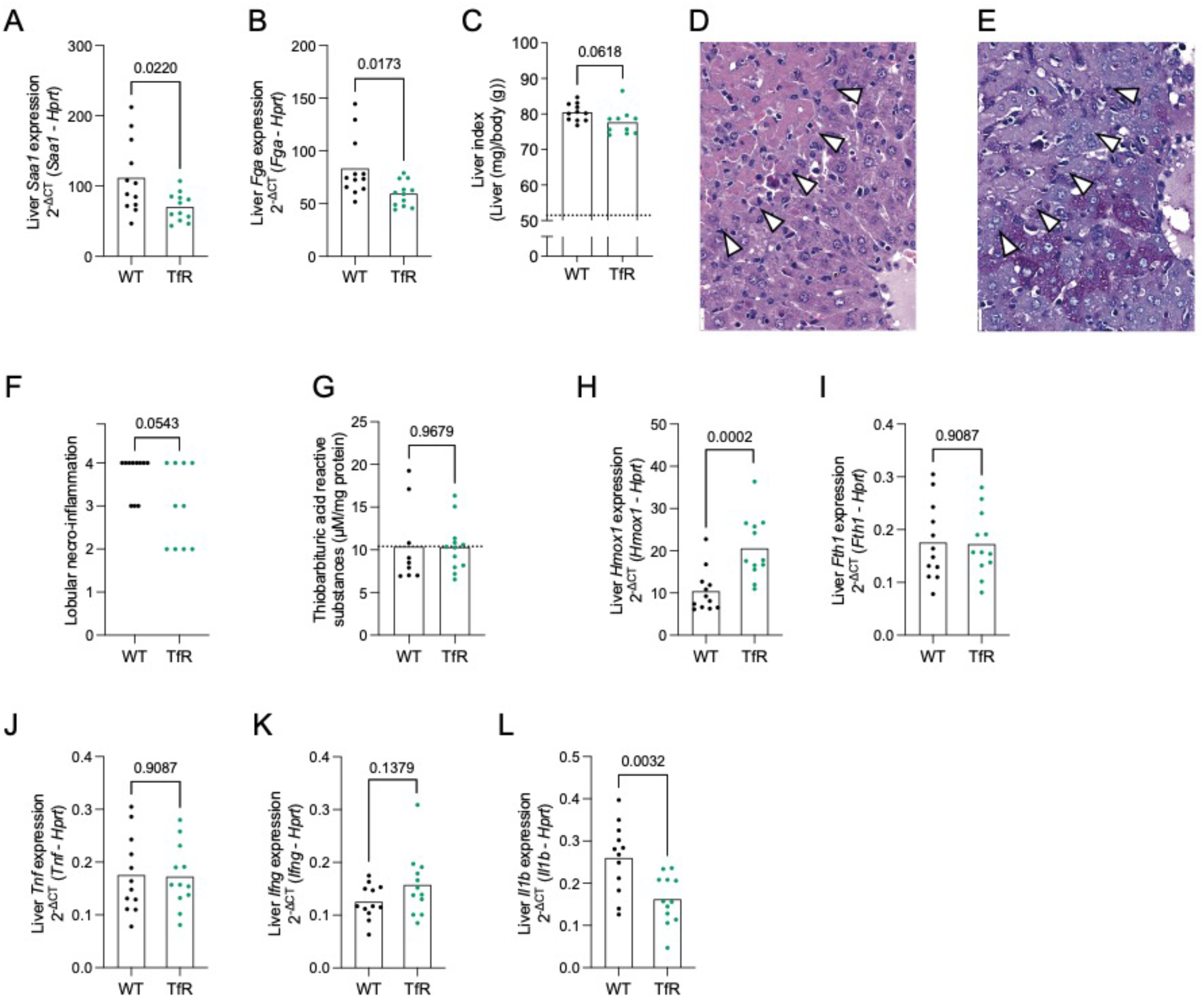
Decreased cellular iron uptake attenuates *P. chabaudi* induced liver damage. Hepatic response of *P. chabaudi* infected C57BL/6 (WT) and *Tfrc^Y20H/Y20H^* (TfR) mice, 8 dpi. **A-B)** Liver gene expression of *Saa1* (A) and *Fga* (B) of *P. chabaudi* infected WT and TfR mice. Mean, Welch’s t-test, n=12. **C)** Liver index of *P. chabaudi* infected WT and TfR mice. Mean, Welch’s t-test, n=10-11. **D-E)** Higher magnification depiction of H&E (D) and PAS (E) stained liver sections from a representative *P. chabaudi* infected WT mouse. The arrowheads indicate areas of confluent necrosis, featuring lobular disarray, lympho-histiocytic inflammation, acidophil body formation, and glycogen depletion. Original magnification 200×, scale bar 20µm. **F)** Blinded scoring of lobular necro-inflammatory activity. Mann-Whitney test, n=10-11. **G)** Hepatic malondialdehyde (MDA), quantified as an indirect measurement or ROS, using a thiobarbituric acid reactive substances assay in *P. chabaudi* infected WT and TfR mice. Mean, Welch’s t-test, n=10-12. **H-J)** Liver gene expression of *Tnf* (H), *Ifn* (I) and *Il1b* (J). Mean, Welch’s t-test on untransformed (H&J) or log transformed data (I), n=12.

## Notes

### Competing Interest Statement

The authors have declared no competing interest.

### Summary of Updates

The manuscript has been revised according to reviewers comments received through Review Commons.

